# Sterile- or pathogen-induced endolysosomal damage activate the NLRP6 inflammasome in human intestinal epithelial cells

**DOI:** 10.1101/2025.01.23.634286

**Authors:** Alexandra Boegli, Elliott M. Bernard, Louise Lacante, Gaël Majeux, Vanessa Mack, Petr Broz

## Abstract

NLRP6 controls host defense against bacteria and viruses in the gastrointestinal tract by a poorly understood mechanism. Here, we report that NLRP6 forms an inflammasome upon endolysosomal damage caused by sterile triggers or bacterial pathogens such as *Listeria monocytogenes* in human intestinal epithelial cells (IECs). NLRP6 activation required Listeriolysin O-dependent cytosolic invasion of *L. monocytogenes* and drove IEC pyroptosis via ASC/Caspase-1-mediated GSDMD cleavage. NLRP6 inflammasome formation was independent of bacterial PAMPs, such as lipoteichoic acid or dsRNA, which were previously reported to activate NLRP6. *L. monocytogenes* mutants deficient in cell-to-cell spread or escape from secondary vacuoles induced less cell death, linking bacteria-induced endolysosomal damage to NLRP6 activation. Finally, sterile endolysosomal damage recapitulated pathogen-induced NLRP6 activation and drove IEC pyroptosis. In summary, our study reveals that NLRP6 allows IECs to detect endolysosomal damage, thereby not only making it responsive to pathogens but more generally to wide-ranging sources of pathological endolysosomal damage.

## Introduction

The human body constantly encounters pathogens at barrier tissues such as the skin, lung and gastro-intestinal tract. First line defenses like mucus secretion and antimicrobial peptides, but also specialized immune cells, like macrophages, protect barrier tissues from being invaded by these pathogens. To avoid these extracellular defenses, many pathogens have developed strategies to invade and replicate within epithelial cell layers, and even to spread intracellularly from cell-to-cell. One such pathogen is *L. monocytogenes,* a foodborne, Gram-positive bacterium that can cause severe diseases such as meningitis and sepsis in susceptible patients ^1,2^. *Listeria* enters enterocytes in the gastro-intestinal tract and escapes from the endolysosomal system using Listeriolysin O (LLO), a pore-forming toxin, to damage the endolysosomal membrane ^3^. In the host cell cytosol, *Listeria* replicates and uses the effector ActA, which recruits host Arp2/3, to polymerize actin to move within the infected cell and to spread to neighboring cells ^4^. This strategy allows the bacteria to avoid resurfacing to the hostile extracellular compartment and can even promote the dissemination to distal organs via migrating cells such as macrophages ^5^. Other enteric bacterial pathogens like *Shigella flexneri* also use actin-based motility to move from cell to cell and can cause high morbidity and mortality in risk groups ^6,7^.

To sense intracellular pathogens, host cells express pattern recognition receptors (PRRs) that detect conserved microbial molecules, such as modified nucleic acids, cell wall components, or a disruption of cellular homeostasis caused by toxins or other virulence factors. A subset of cytosolic PRRs, namely PYRIN, AIM2 and members of the NOD-like receptor family (NLRs), induce the formation of multi-protein signaling complexes, known as inflammasomes. Inflammasomes recruit and activate the protease Caspase-1, either directly or via the adaptor ASC, which then cleaves the pro-forms of IL-1β/-18, to produce the mature forms of these cytokines, and the induction of pyroptosis via the cleavage and activation of the pore-forming cell death executioner GSDMD ^8^. Inflammasome-induced cell death is particularly important in barrier tissues, as it can, for example, promote the extrusion of infected cells into the intestinal lumen thereby reducing bacterial loads in the intestinal epithelium, as shown for NLRC4-mediated restriction of the enteric pathogens *S. flexneri* or *Salmonella enterica* serovar Typhimurium ^9–11^.

The NLR family member NLRP6 is mainly expressed in enterocytes of the intestine and plays an important role in regulating susceptibility to bacterial and viral infections ^12–19^. Furthermore, NLRP6 is linked to colorectal cancer and inflammatory bowel disease, as well as neuroinflammation ^20,21^. While some of its functions were proposed to depend on regulating NFϕB or type-I-interferon (IFN) signaling ^14,15^, NLRP6 has also been reported to form an inflammasome and thereby regulate IL-18 release *in vivo*^17,18,22,23^. The NLRP6 inflammasome has also been suggested to regulate autophagy and mucus secretion by Goblet cells, although others have not found a role in baseline mucus layer formation ^13,19^. Several chemically and structurally different ligands were proposed to activate NLRP6. Among these are bacterial metabolites, double stranded viral RNA (dsRNA) and lipoteichoic acid (LTA), a cell wall component of Gram-positive bacteria such as *L. monocytogenes* and *Staphylococcus aureus* ^17,18,23^.

In this study we show that human and mouse NLRP6 form an inflammasome in response to *L. monocytogenes* infection, and that this requires Listeriolysin O-dependent bacterial entry into the host cell cytosol. In *Listeria*-infected human intestinal epithelial cells (IECs), NLRP6 activation resulted in the formation of ASC specks that induced Caspase-1-dependent, but Caspase-4-independent, GSDMD cleavage and pyroptotic cell death.

Functional characterization of NLRP6 identified critical roles for a positively charged patch within the FISNA domain and the Walker A and B motif in the NACHT domain of NLRP6. The generation of chimeric receptors between NLRP6 and the closely related NLRP3 showed that *Listeria* recognition was mediated via the NLRP6 NACHT domain. Further investigating NLRP6 signal recognition, we found that none of the previously reported NLRP6-activating pathogen associated molecular patterns (PAMPs)^17,23^ activated NLRP6 in our system. Indeed, even if we transfected whole bacterial lysate, composed of a mixture of diverse PAMPs NLRP6 did not get activated. In contrast, NLRP6 activation required the presence of live, replicating bacteria in the cytosol, suggesting that NLRP6 does not sense PAMPs but pathogen-induced alterations to cellular homeostasis. We tested the importance of cell-to-cell spread and found that both blocking actin-based motility and escape from double-membrane vacuoles after cell-to-cell spread strongly reduced NLRP6 activation, indicating that vacuolar escape during secondary infections and the resulting endolysosomal damage is a prerequisite for NLRP6 activation. Confirming that NLRP6 acts as a sensor for endolysosomal integrity, we finally found that inducing sterile endolysosomal damage also led to NLRP6 activation and Caspase-1-dependent IEC death, completely independent of any bacterial PAMP. In summary these data show that NLRP6 does not get activated by pathogen-derived danger signals as previously published but responds to the integrity of endolysosomal pathways.

## Results

### NLRP6 detects the entry of *L. monocytogenes* into the host cell cytosol

NLRP6 is predominantly expressed in primary IECs, with no expression found in mouse bone-marrow derived macrophages even after priming with TLR ligands or interferon-γ (IFNγ) ^21,23^ (**Fig. S1A**). Therefore, we developed an NLRP6 inflammasome reconstitution assay in HEK293T cells based on the inflammasome adaptor protein ASC that forms macromolecular inflammasome assemblies referred to as ASC specks. For this assay, HEK293T cells stably expressing GFP- or mCherry-tagged ASC (GFP-/mCherry-ASC^tg (transgenic)^ were transiently transfected with an NLRP6 expression plasmid. Since NLRs are prone to auto-activate when overexpressed, we first determined the dynamic range of the assay by transfecting increasing amounts of plasmid DNA encoding human NLRP6. We found that with increasing amounts of transfected plasmid the percentage of ASC speck-positive cells reached a plateau at 50-60% of the total population as determined by flow cytometry-based quantification of ASC speck formation ^24^ (**Fig. S1B**).

To probe for signal-specific NLRP6 activation, we chose a concentration of human NLRP6 vector DNA that caused only basal ASC-speck formation upon transfection into GFP-ASC^tg^ HEK293T (80 ng/300,000 cells). These cells were infected with two strains of the Gram-positive bacteria *L. monocytogenes*, as well as *S. aureus*, which were proposed to induce NLRP6 activation and the Gram-negative bacterium *S.* Typhimurium as a control ^15–17^. While *S. aureus* and *S.* Typhimurium did not cause elevated levels of ASC speck formation, *L. monocytogenes* infection resulted in up to 40% of ASC speck-positive cells (**Fig. 1A, B and S1C**), indicating that *Listeria* infection triggered NLRP6 activation. To validate the specificity of this response we expressed NLRP6^W53E^, which is defective for ASC recruitment ^25^, and found that the mutant protein did not allow ASC speck formation after *Listeria* infection (**Fig. 1B**). Additionally, we also repeated the assays with murine NLRP6, which shares 71.14% amino acid identity with human NLRP6 (BLASTP 2.16.0+), and infected these cells with *L*. *monocytogenes*, this also induced ASC speck formation, indicating that NLRP6 activation upon *Listeria* infection is conserved between mouse and human homologs (**Fig. 1C, D and S1D**).

**Figure 1.**
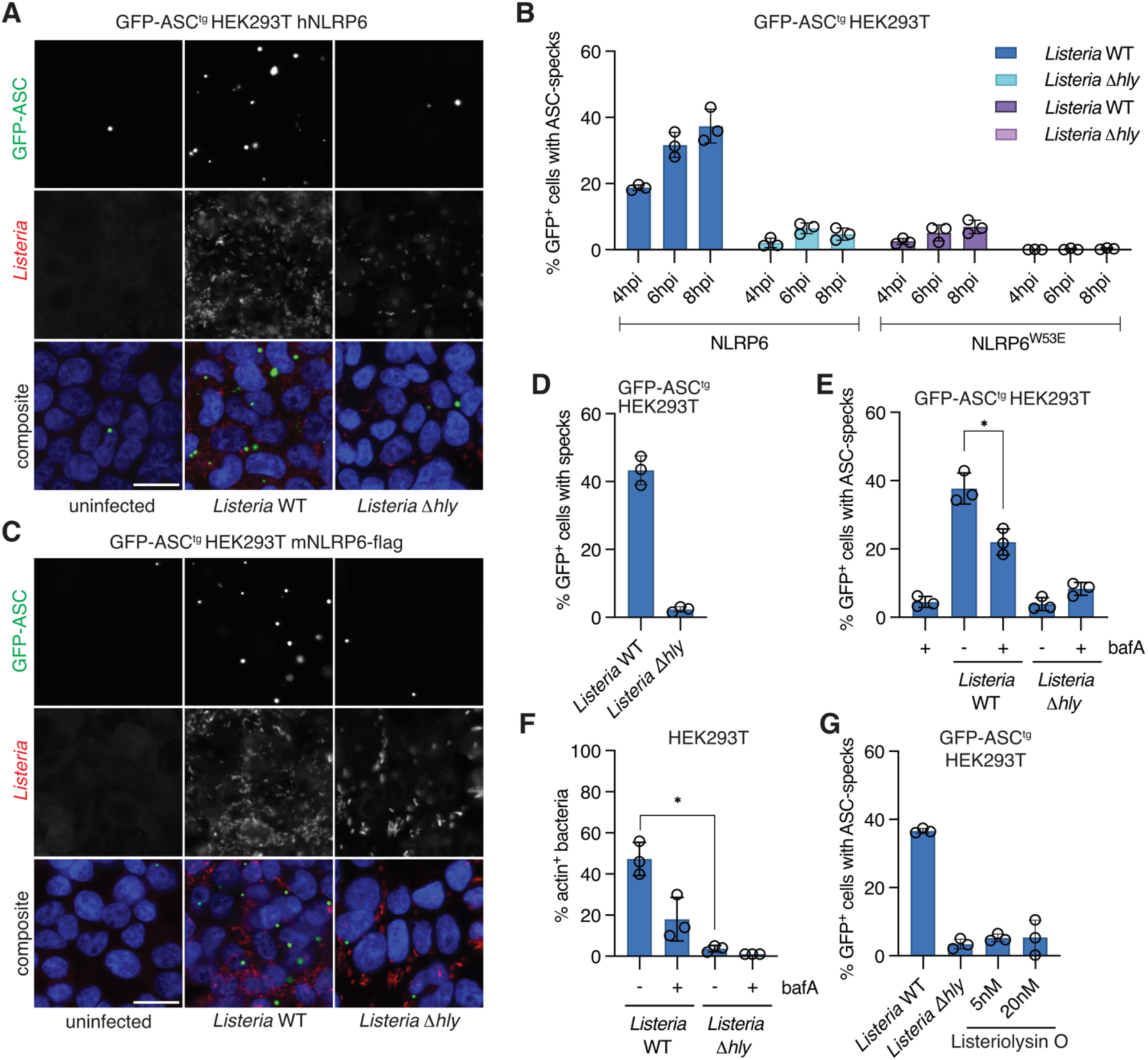
NLRP6 detects the entry of *Listeria* into the host cell cytosol. **A.** Representative micrographs of hNLRP6-expressing GFP-ASC^tg^ HEK293T cells infected with WT or τ<*hly L. monocytogenes* EGD for 6h at MOI 20. **B.** Flow-cytometry based quantification of ASC speck formation in hNLRP6 or hNLRP6^W53E^-expressing GFP-ASC^tg^ HEK293T cells infected with WT or τ<*hly L. monocytogenes* EGD for 4-8h. hpi: hours post infection. **C-D.** Representative micrographs and quantification of ASC speck formation by flow cytometry in mNLRP6-expressing GFP-ASC^tg^ HEK293T cells infected with WT or τ<*hly L. monocytogenes* EGD for 6h **E.** ASC speck formation in hNLRP6-expressing GFP-ASC^tg^ HEK293T cells left untreated or treated with bafilomycin A1 (bafA) for 2h and infected with WT or τ<*hly L. monocytogenes* EGD for 6h **F.** Percentage of actin positive, intracellular *Listeria* after 1h of infection in HEK293T cells treated as in E. **G.** ASC speck formation of hNLRP6-expressing GFP-ASC^tg^ HEK293T cells infected with WT or τ<*hly L. monocytogenes* EGD for 6h at MOI 20 or treated with 5 or 20nM purified Listeriolysin O for 6h. Graphs show mean ± SD from three pooled independent experiments. Quantification in panel F was performed on maximum projection confocal micrographs (Fig. S1E) of ten fields of view and at least 600 bacteria per experiment and condition. Each data point represents the mean of one experiment. Images are representative of at least three independent experiments. Scale bar represents 20μm. *P < 0.05 (unpaired t-test).

*L. monocytogenes* employs the pore forming toxin Listeriolysin O to escape from the endolysosomal compartment into the cytosol ^3^. We found that wild-type bacteria caused progressive NLRP6-dependent ASC speck formation over time, while the *τιhly* strain, which lacks Listeriolysin O, caused no significant NLRP6 activation (**Fig. 1A-B**). Optimal activity of Listeriolysin O requires the low pH of the phagolysosome, allowing the bacteria to escape this compartment without causing excessive damage to other host cell membranes in the process ^26–28^. Consistently, preventing the acidification of endolysosomes with bafilomycin A1 efficiently reduced both the escape of wild-type *Listeria* into the cytosol, as judged by reduced numbers of actin-positive bacteria, and NLRP6-dependent ASC speck formation (**Fig. 1E-F, S1E**). We also tested if Listeriolysin O itself could trigger NLRP6 activation. At the concentrations used, purified Listeriolysin O caused plasma membrane permeabilization (**Fig. S1F**) but did not trigger NLRP6-dependent ASC-speck formation (**Fig. 1G**), indicating that Listeriolysin O promoted NLRP6 activation indirectly by allowing cytosolic entry of *Listeria*. Furthermore, while *Listeria* infection caused NLRP6 activation, it did not induce host cell permeabilization, thus excluding plasma membrane permeabilization as a potential trigger for NLRP6 activation (**Fig. S1G**). In summary, these experiments showed that the entry of *L. monocytogenes* into the host cell cytosol is required for the activation of NLRP6 in HEK293 cells.

### NLRP6 induces Caspase-1-dependent GSDMD cleavage and pyroptosis in infected human IECs

We next wanted to study NLRP6 activation by *Listeria* infection in a more physiological setting. As HEK293T cells lack critical inflammasome components such as Caspase-1 and GSDMD, we turned to HIEC-6 (human intestinal epithelial cell-6) enterocytes, derived from human fetal small intestine ^29^, to determine if NLRP6 induces the full inflammasome pathway upon activation. Immunoblotting confirmed that HIEC-6 express the inflammasome components ASC, Caspase-1, Caspase-4, and GSDMD, but do not endogenously express NLRP6 (**Fig. S2A**) or NLRP3 (**Fig. S2B**). We therefore reconstituted these cells with doxycycline-inducible wild-type NLRP6 or the NLRP6^W53E^ mutant (**Fig. S2A**) before infecting them with *L. monocytogenes*. Consistent with a specific activation of NLRP6, we observed that wild-type *L. monocytogenes* caused significantly elevated levels of ASC speck formation, GSDMD processing, and LDH release (a measure for lytic cell death) in HIEC-6 treated with doxycycline to induce NLRP6^WT^ expression but not in non-doxycycline treated, and thus non-NLRP6 expressing, controls (**Fig. 2A-D**). Of note, doxycycline treatment induced some background inflammasome activation due to NLRP6 autoactivation, as visible by background ASC-speck formation and GSDMD processing. Consistent with data obtained from HEK293T cells, we found that the interaction of NLRP6^PYD^ with ASC^PYD^ was required for inflammasome assembly in HIEC-6 cells, since expression of NLRP6^W53E^ did not allow ASC speck formation, GSDMD cleavage or LDH release after wild-type *L. monocytogenes* infection (**Fig. 2A-D**). In addition, we observed that NLRP6 was recruited to ASC specks, further indicating that NLRP6 is the driver of inflammasome responses in these cells (**Fig. 2A**). As observed in HEK293T cells, the comparison of wild-type and τι*hly L. monocytogenes* infections showed that cytosolic entry was required for NLRP6-dependent ASC speck formation, GSDMD cleavage and cell death in NLRP6^tg^ HIEC-6 cells (**Fig. 2A-D**). To determine the kinetics of *Listeria*-induced NLRP6 activation in HIEC-6 cells, we next assayed propidium iodide (PI) uptake, a measure of plasma membrane permeabilization by GSDMD pores, in cells infected with wild-type and τι*hly L. monocytogenes.* This analysis showed that the PI signal started to increase in cells infected with WT bacteria compared to cells infected with τι*hly* bacteria or cells not expressing NLRP6 at approximately 3 h post infection (**Fig. 2E**), indicating that NLRP6 activation occurs within few hours after bacterial escape into the cytosol.

**Figure 2.**
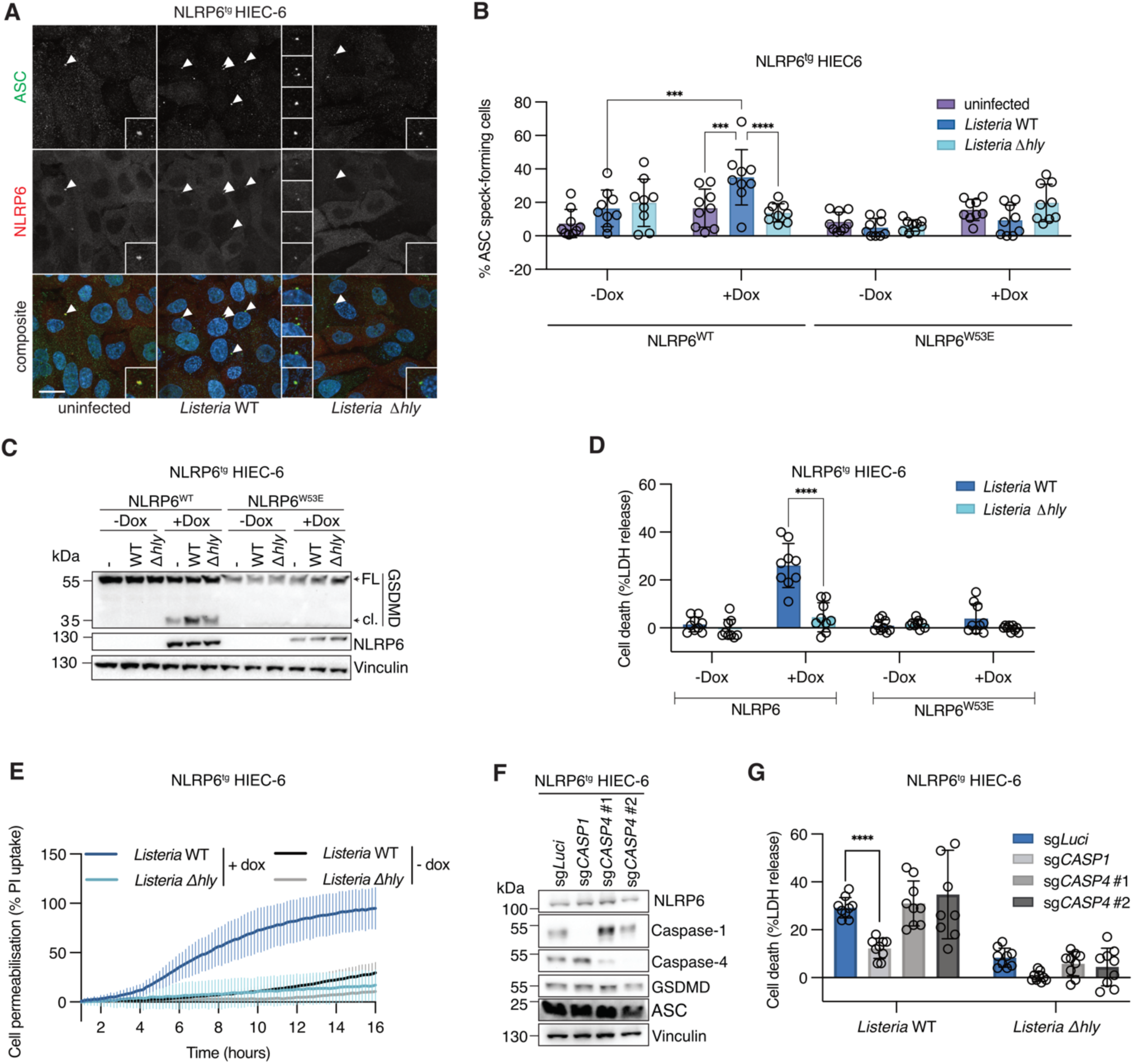
NLRP6 induces Caspase-1-dependent GSDMD cleavage and pyroptosis in *Listeria* infected human IECs. **A-B.** Representative maximum projection confocal micrographs and ASC speck quantification in NLRP6-WT^tg^ or NLRP6-W53E^tg^ HIEC-6 cells induced or not with 1μg/ml doxycycline overnight and infected with WT or τ<*hly L. monocytogenes* EGD for 8h. Arrowheads indicate regions in insets. **C-D.** GSDMD processing and LDH release from NLRP6-WT^tg^ or NLRP6-W53E^tg^ HIEC-6 cells induced or not with 1μg/ml doxycycline overnight and infected with WT or τ<*hly L. monocytogenes* EGD for 8h. **E.** Time course of propidium iodide (PI) uptake by NLRP6-WT^tg^ HIEC-6 cells induced or not with 1μg/ml doxycycline overnight and infected with WT or τ<*hly L. monocytogenes* EGD. **F-G.** Protein expression and LDH release from NLRP6-WT^tg^ HIEC-6 control cells (sg*Luci*) or polyclonal populations lacking Caspase-1 (sg*CASP1*) or Caspase-4 (sg*CASP4* #1 and #2) induced with 1μg/ml doxycycline overnight and infected with WT or τ<*hly L. monocytogenes* EGD for 8h. Graph B shows mean ± SD from three fields of view each for three pooled experiments with at least 200 cells analyzed per condition per experiment. Each data point represents one field of view. Graphs D, E and G show mean ± SD from three pooled experiments with three technical replicates each. Images and blots are representative of three independent experiments. FL (full length), cl. (cleaved). Scale bar represents 20μm. *P < 0.05, **P < 0.01, ***P < 0.001, ****P < 0.0001 (two-way ANOVA with Šídák’s multiple comparisons).

A previous study proposed that in mouse macrophages transfection of LTA of *S. aureus* induces NLRP6- and ASC-dependent Caspase-11 activation, which then promotes Caspase-1 activation and interleukin-1β (IL-1β)/IL-18 maturation without the induction of pyroptosis ^17^. To study the contribution of Caspase-1 and Caspase-4 (the human ortholog of murine Caspase-11), we used CRISPR/Cas9 to inactivate these genes in NLRP6^tg^ HIEC-6 cells (**Fig. 2F**) and infected the cells with *L. monocytogenes.* While deletion of Caspase-1 significantly reduced pyroptotic cell death as measured by LDH release, Caspase-4 deficiency had no impact on NLRP6-induced death (**Fig. 2G**). In summary, our data show that NLRP6 is activated upon cytosolic entry of *L. monocytogenes* and consequently polymerizes ASC, leading to Caspase-1 recruitment, GSDMD cleavage and pyroptotic cell death in human IECs without any contribution of the non-canonical Caspase-4 inflammasome pathway. Furthermore, the results confirm that the Listeriolysin O-dependent escape of *Listeria* from the endolysosomal compartment is an essential prerequisite for NLRP6 activation in human epithelial cells.

### NLRP6 shares structural similarities and ligands with NLRP3

To understand how NLRP6 detects *Listeria* infection, we next characterized its mode of activation, and the domain(s) implicated in this process. Sequence alignment showed that NLRP6 is highly similar to NLRP3, comprising an N-terminal PYD domain, followed by a linker, a NACHT domain and C-terminal LRRs (**Fig. 3A**). NLRP3 has been proposed to detect alterations of cellular homeostasis, such as the disruption of the TGN or endosomal trafficking, but the exact nature of the signal remains unknown. Interestingly, NLRP3 was also reported to detect infection by various pathogens, also including *L. monocytogenes* ^30,31^. Given these similarities, we tested if NLRP3 and -6 share additional activators by infecting NLRP3- or NLRP6-expressing ASC-GFP^tg^ HEK293T with wild-type or *τιhly L. monocytogenes*, or treating them with nigericin and imiquimod, two well-known NLRP3 activators. Wild-type *L. monocytogenes* induced robust ASC-speck formation in both NLRP3 and NLRP6 expressing cells, which was dependent on cytosolic entry in both instances. By contrast, while nigericin and imiquimod efficiently triggered NLRP3 activation, they failed to activate NLRP6 (**Fig. 3B**), revealing that NLRP6 does not share all activators with NLRP3. Since potassium efflux has previously been shown to be required for NLRP3 activation by nigericin and other triggers, we tested its impact on NLRP6 activation but found that addition of extracellular potassium, to prevent potassium efflux, did not reduce *Listeria*-induced NLRP6 activation (**Fig. 3B**). Interestingly, extracellular potassium also did not block *Listeria*-induced NLRP3 activation, indicating a mode of activation distinct from nigericin (**Fig. 3B**).

**Figure 3.**
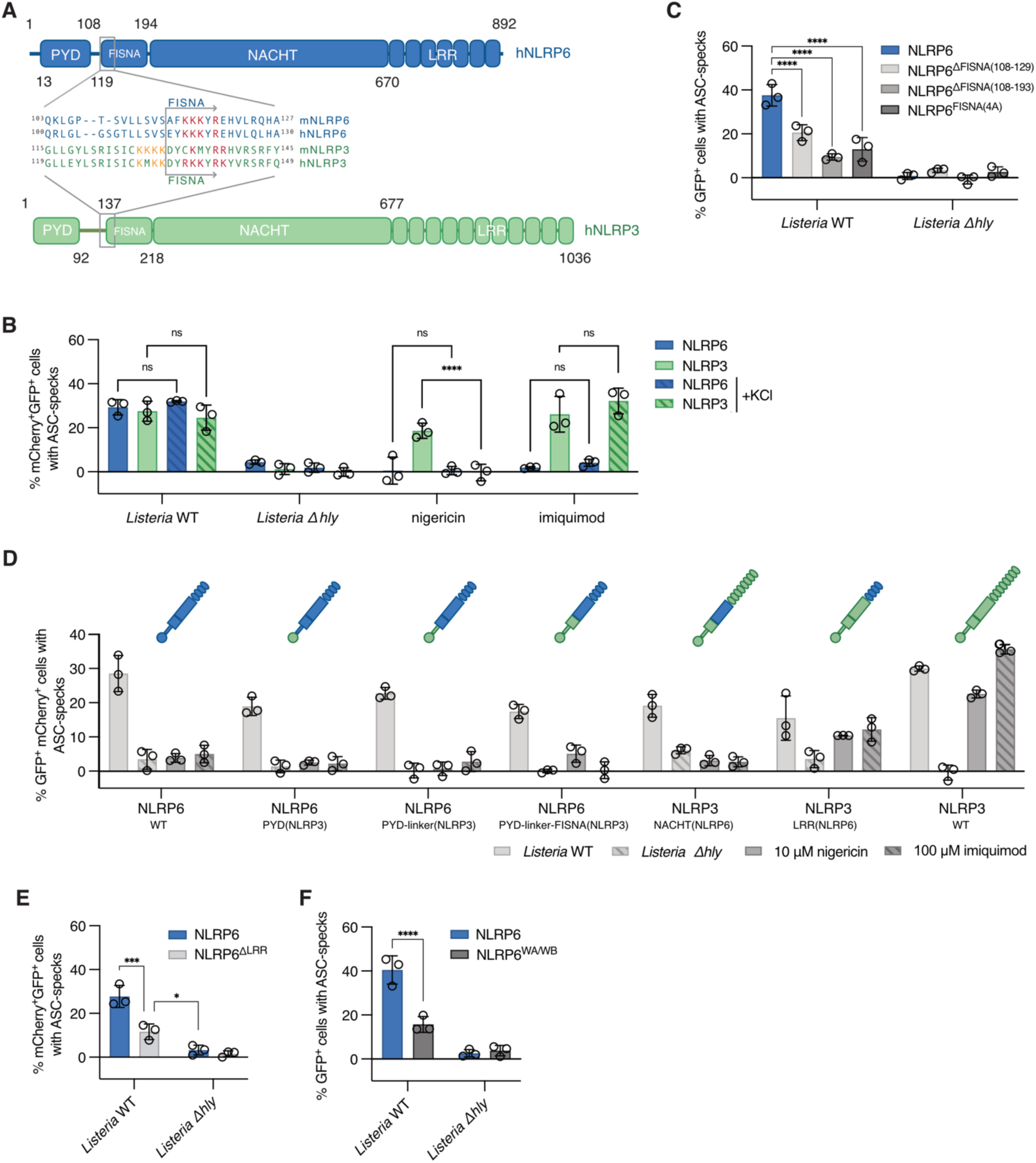
Molecular characterization of the NLRP6 receptor. **A.** Schematic representation of human NLRP6 and NLRP3 showing their domain structures and the respective polybasic regions within the FISNA of mouse and human NLRP3 and 6 aligned. **B.** Flow cytometry-based quantification of ASC speck formation in GFP-ASC^tg^ HEK293T cells expressing hNLRP6-mCherry or hNLRP3-mCherry infected with WT or τ<*hly L. monocytogenes* EGD for 6h, treated with nigericin for 1h or with imiquimod for 6h in presence or absence of 60 mM extracellular KCl. **C.** Flow cytometry based quantification of ASC speck formation in GFP-ASC^tg^ HEK293T cells, expressing the indicated NLRP6 proteins, infected with WT or τ<*hly L. monocytogenes* EGD for 6h **D.** Flow cytometry based quantification of ASC speck formation in GFP-ASC^tg^ HEK293T cells expressing either NLRP6-WT, NLRP3-WT or the indicated chimeric NLRP6-3 proteins with mCherry tags, infected with WT or τ<*hly L. monocytogenes* EGD for 6h, treated with nigericin for 1h or with imiquimod for 6h. **E.** Flow cytometry based quantification of ASC speck formation in GFP-ASC^tg^ HEK293T cells, expressing NLRP6^τιLRR^-mCherry, infected with WT or τ<*hly L. monocytogenes* EGD for 6h. **F.** Flow cytometry based quantification of ASC speck formation in GFP-ASC^tg^ HEK293T cells, expressing NLRP6^WA/WB^, infected with WT or τ<*hly L. monocytogenes* EGD for 6h. Graphs show mean ± SD from three pooled experiments. ns = non-significant, **P < 0.01; ***P < 0.001, ****P < 0.0001 (two-way ANOVA with either Tukey’s multiple comparisons test (B), Dunett’s multiple comparisons test (C,E) or Uncorrected Fisher’s LSD (F)).

A characteristic feature of NLRP3 is its FISNA domain, which is located between the linker and NACHT domain. Previous work has suggested that nigericin and imiquimod sensing maps to the linker-FISNA-domains of NLRP3 and recent structural studies showed that several disordered regions in the NLRP3-FISNA, such as loop1 and middle region/loop 3 (**Fig. S3A**), fold during the transition from the inactive to the active receptor, thus allowing receptor oligomerization and correct positioning of the PYD ^32,33^. Therefore, mutations within the FISNA domain could critically affect these conformational changes that are essential for receptor activation, rather than signal recognition/sensing. NLRP6 features a conserved polybasic motif preceding the NACHT in a domain that shows sequence similarity to the FISNA of NLRP3 and NLRP12 and which we thus propose to define as the NLRP6 FISNA (**Fig. 3A**) ^32,34^. Moreover, pairwise alignments of the predicted structure of the NLRP6-FISNA (Alphafold v2) with inactive and active NLRP3, shows that it is proposed to adopt a similar structure with helix 1, loop1, loop 2 and loop3, which likely would fold into helix 2 during receptor activation (**Fig. S3A**). To test the role of the FISNA in NLRP6 activation, we expressed two deletion mutants (aa108-129 and aa108-193) in GFP^tg^ HEK293T and found that both deletions rendered the receptor unable to induce ASC speck formation upon *Listeria* infection (**Fig. 3C**). The larger FISNA deletion (aa108-193) showed partially impaired auto-oligomerization, potentially indicating a structural defect that reduced ASC recruitment (**Fig. S3B-C**).

Mouse NLRP3 is activated by the recruitment to phosphatidylinositol-4-phosphate (PI4P)-enriched membranes via a positively charged motif (KKKK^130^) in the linker domain ^35^. Mutation of the corresponding motif (KMKK^134^) in human NLRP3 has been shown to have no impact on activation, potentially due to redundancies with a second polybasic motif located within the FISNA of human NLRP3 (RKKYRK^142^) ^32^. While NLRP6 does not feature the first polybasic motif, it features the second polybasic motif in its FISNA (KKKYR^123^) (**Fig. 3A**). Mutating these four positively charged residues to alanine led to a significant decrease in NLRP6-dependent ASC-speck formation upon *Listeria* infection, even though the receptor was still able to generate ASC specks through auto-activation upon high levels of overexpression (**Fig. 3C, S3B-C**). Similarly, NLRP3^FISNA(5A)^ was significantly impaired in recruiting ASC in response to *Listeria* infection, nigericin and imiquimod treatment, but this receptor also showed impaired auto-oligomerization upon overexpression (**Fig. S3D-F**).

In summary, these data show that even though NLRP3 and NLRP6 activation depends on conserved structural motifs, such as the FISNA domain and its polybasic motifs, the two inflammasome receptors recognize a distinct set of ligands. Since the NLRP3 linker-FISNA is implicated in the spatial organization of NLRP3 in both the inactive cage-oligomer and the inflammasome disc assembly, it is likely that the NLRP6 FISNA plays a similar role in the formation of active NLRP6 complexes ^33,36^. However, a function for the FISNA domain in signal sensing cannot be excluded from these findings.

### The NLRP6 NACHT plays an important role in receptor activation

To identify which domains within NLRP6 sense *Listeria* infection, we created a set of chimeras between NLRP6 and NLRP3, using corresponding domains in NLRP3 as place holders to generate functional receptors, and tested their responsiveness to *Listeria* infection in ASC-GFP^tg^ HEK293T cells. Apart from a few receptors, most chimeras were expressed to the same level as wild-type NLRP3 or NLRP6 and caused similar levels of ASC speck formation upon high levels of overexpression (**Fig. S3G-H**). We next tested signal specific activation and found that exchanging the PYD, the PYD-linker or the PYD-linker-FISNA region of NLRP6 with the corresponding domains from NLRP3 had no significant impact on *Listeria*-induced ASC speck formation, nor did it confer a response to the NLRP3 activators nigericin or imiquimod (**Fig. 3D**). This indicated that while the NLRP6 PYD-linker-FISNA region is important for ASC speck formation by mediating receptor oligomerization and the recruitment of ASC, it likely does not determine signal specificity. Therefore, *Listeria* recognition by NLRP6 must rely on either the NACHT or LRR domains, or both together. Consequently, our data also imply that NLRP3 signal specificity for nigericin and imiquimod is predominantly conferred by its NACHT-LRR region, and not the PYD-linker-FISNA region.

To determine whether the NACHT domain of NLRP6 was sufficient to confer NLRP6 signal specificity onto NLRP3, we replaced the NACHT of NLRP3 with the corresponding domain from NLRP6. The resulting NLRP3^NACHT(NLRP6)^ chimera reacted to *Listeria* infection but was insensitive to nigericin and imiquimod treatment, thus mimicking the signal specificity profile of NLRP6 (**Fig. 3D**). This result demonstrated that the NLRP6 NACHT could confer NLRP6-like ligand specificity on NLRP3, indicating that in both NLRP6 and NLRP3, specific ligand detection requires their respective NACHT domains.

To investigate if the NLRP6 LRRs also play a role in sensing of an activation signal, we generated the NLRP3^LRR(NLRP6)^ receptor chimera. This protein showed ASC speck induction upon *Listeria* infection but also responded to nigericin and imiquimod treatment as observed with NLRP3 wild type (**Fig. 3D**). Thus, NLRP6 LRRs can functionally substitute for NLRP3 LRRs, but do not confer NLRP6-like ligand specificity. Since this indicated that the LRRs mainly serve a structural function that enhances receptor activation, we also deleted the LRRs in NLRP6. The resulting protein showed reduced responses to wild-type *Listeria* infection, yet responses were still higher than the control condition (*τιhly Listeria* infection) and the receptor was fully able to auto-activate (**Fig. 3E, S3I-J**). Thus, we conclude that the NACHT domain is the cognate sensor domain of NLRP6 with an important contribution of the LRRs for full inflammasome activation.

Finally, a hallmark of NLR activation is an ATP-dependent receptor oligomerization into wheel-shaped assemblies, requiring Walker A and B motifs in the NACHT that mediate ATP binding and hydrolysis. NLRs with mutations in either the Walker A or B motif are strongly reduced in their ability to respond to activators, but can oligomerize independently of ATP and recruit ASC upon overexpression ^37–40^ (**Fig. S3K-L**). Mutating the Walker A/B motifs in NLRP6 significantly reduced *Listeria*-induced ASC speck formation, indicating that ATP binding and likely the following hydrolysis is an essential step in NLRP6 activation upon signal specific activation of the receptor (**Fig. 3F**). In conclusion, our data suggest that during *Listeria* infection, NLRP6 ligand sensing mainly maps to its NACHT domain and that ATP binding and hydrolysis are an important prerequisite for NLRP6 inflammasome formation. This implies that NLRP6 functions similarly to NLRC4 and NLRP3, where ATP-dependent receptor oligomerization into wheel-shaped inflammasome assemblies is a prerequisite for ASC speck formation ^37–39,41^.

### NLRP6 activation does not require bacterial PAMPs, but viable cytosolic bacteria

NLRP6 has been proposed to detect several chemically distinct ligands (LTA, RNA or metabolites), all of which could theoretically be released by cytosolic *Listeria* ^18,23,37^. To define if one or several of these were detected by NLRP6, we individually evaluated their ability to activate NLRP6 in ASC-GFP^tg^ HEK293T cells. Even though *Listeria* triggered NLRP6-dependent ASC speck formation (**Fig. 1A-B**), we found that transfection of the Gram-positive cell wall component LTA did not cause any ASC speck formation (**Fig. 4A**), in contrast to published work in primary mouse macrophages ^17^. Double-stranded (ds) RNA has been shown to act as an NLRP6 ligand *in vitro* and to induce NLRP6 clustering and potentially inflammasome formation by liquid-liquid phase-separation^14,23^. However, neither high nor low molecular weight poly(I:C) (a synthetic dsRNA analogue) induced NLRP6-dependent ASC-speck formation upon transfection, even at high concentrations (**Fig. 4A**, 10 μg/300,000 cells). To verify that LTA and poly(I:C) was indeed delivered into cells by transfection, we transfected BODIPY-labelled LTA or fluorescein-labelled poly(I:C) respectively. Additionally, we transfected glycine quenched BODIPY and fluorescein to control for unspecific colocalization. While efficient delivery of LTA-BODIPY and poly(I:C)-fluorescein was detected, we found low to no colocalization with NLRP6, consistent with the inability of LTA or poly(I:C) to activate NLRP6 and induce ASC speck formation (**Fig. 4B, S4A-B**). HEK293T cells might lack a co-factor needed for LTA- or dsRNA-induced NLRP6 activation, such as DHX15 that has been proposed to mediate NLRP6-induced IFN responses ^23^. Therefore, we tested LTA and poly(I:C) transfection in the HIEC-6 model, either inducing NLRP6 expression with doxycycline or not. LTA, as well as Lipofectamine by itself, did not induce cell death as measured by LDH release, while poly(I:C) transfection did cause cell death, but independently of NLRP6 expression (**Fig. 4D**). We thus conclude that neither LTA nor poly(I:C) are ligands of NLRP6 that are capable of activating NLRP6 in our model systems.

**Figure 4.**
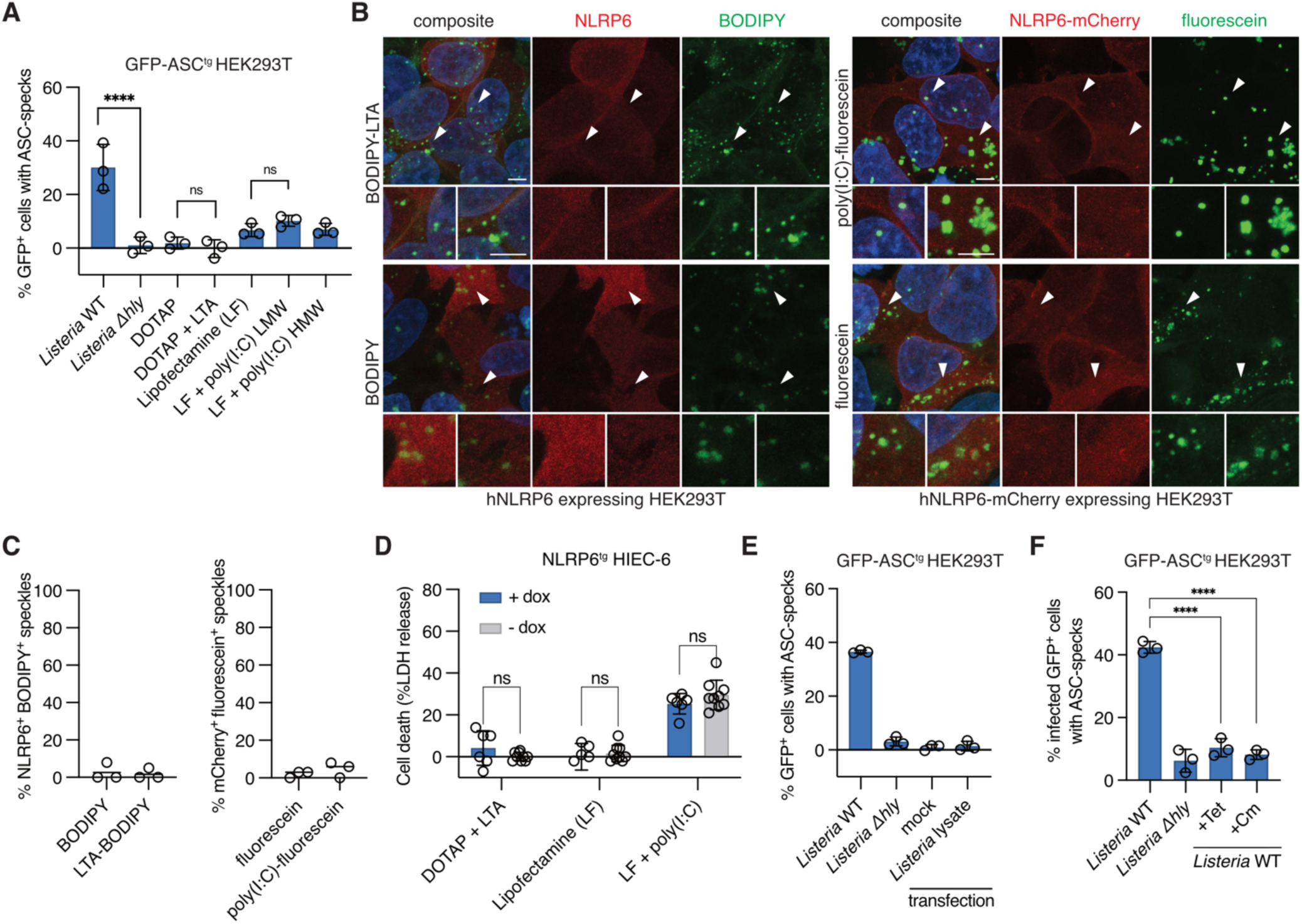
NLRP6 activation does not require bacterial PAMPs, but viable cytosolic bacteria. **A.** Flow cytometry-based quantification of ASC speck formation in hNLRP6-expressing GFP-ASC^tg^ HEK293T cells either infected with WT or τ<*hly L. monocytogenes* EGD for 6h or mock transfected with DOTAP or Lipofectamine (LF) only or transfected with LTA, poly(I:C) LMW or HMW for 24h at 10μg/300,000 cells. **B.** Representative maximum projection confocal micrographs after transfection of hNLRP6 or hNLRP6-mCherry expressing HEK293T cells with LTA-BODIPY (7.5μg/300,000), equivalent volume BODIPY only, poly(I:C)-fluorescein (250ng/300,000 cells) or equivalent volume fluorescein only for 6h. Arrowheads indicate regions in insets. **C.** Quantification of images represented in B. For each condition, ten fields of view and at least 800 BODIPY or fluorescein speckles in NLRP6 positive cells were analyzed for NLRP6 colocalization per experiment. Three independent experiments were performed. **D.** LDH release from NLRP6-WT^tg^ HIEC-6 cells induced or not with 1μg/ml doxycycline overnight and transfected with Lipofectamine (LF) only or transfected with LTA or poly(I:C) for 8h. **E.** Flow cytometry-based quantification of ASC speck formation in hNLRP6-expressing GFP-ASC^tg^ HEK293T cells either infected with WT or τ<*hly L. monocytogenes* EGD, mock transfected or transfected with *L. monocytogenes* EGD lysates for 6h. **F.** Flow cytometry based quantification of ASC speck formation in hNLRP6-expressing GFP-ASC^tg^ HEK293T cells infected with WT or τ<*hly L. monocytogenes* for 6h, in the presence of Tetracycline (Tet) or Chloramphenicol (Cm) added after 45min of infection. Graphs show mean ± SD from three pooled experiments. Images are representative of three independent experiments. Scale bars represent 5μm. ns = non-significant, ****P < 0.0001 (one-way ANOVA with either Tukey’s multiple comparisons test (A) or Dunnett’s multiple comparisons test (F), two-way ANOVA with Tukey’s multiple comparisons test (D)).

Since we could not confirm NLRP6 activation by these two published ligands, we investigated if other PAMPs that are known to be released by *Listeria* caused NLRP6 activation. Intracellular *Listeria* passively release the secondary messenger c-di-AMP, which is bound by STING to induce type I IFN ^42^. We thus infected GFP-ASC^tg^ HEK293T cells expressing NLRP6 with *Listeria* mutants releasing more (τι*mdrMCAT*) or less (tetR::tn) c-di-AMP due to altered efflux pump activity but found similar levels of ASC-speck formation as with wild-type *Listeria* (**Fig. S4C**). Moreover, cytosolic delivery of c-di-AMP by transfection did not induce ASC speck formation (**Fig. S4D**).

Finally, we tested if a yet unknown *Listeria* ligand triggered NLRP6 activation by transfecting whole bacterial lysates, prepared by sonication, into NLRP6-expressing ASC-GFP^tg^ HEK293T cells. However, even the transfection of *Listeria* lysates, which contain a mixture of diverse PAMPs produced by *Listeria*, failed to induce ASC speck formation (**Fig. 4E**). Since these findings implied that NLRP6 does not sense a classical PAMP, we wondered if the presence of live and replicating bacteria in the cytosol was required for NLRP6 activation. We thus treated *Listeria*-infected NLRP6-expressing GFP-ASC^tg^ cells with the bacteriostatic antibiotics tetracycline and chloramphenicol 45 minutes after infection to allow primary escape of the invasion vacuole, but no continuation of the life cycle in the cytosol. Antibiotic treatment both significantly reduced bacterial replication in the cytosol and NLRP6-dependent ASC speck formation in infected cells (**Fig. 4F, S4E**). We thus concluded that primary invasion of the cell and escape into the cytoplasm was insufficient to activate NLRP6 to the full extent, and that, live replicating bacteria are needed for full inflammasome activation.

### NLRP6 recognizes bacterial cell-to-cell spread

Since our data indicated that live, replicating *Listeria* are required for NLRP6 activation, we next investigated if a disturbance of cellular homeostasis caused by *Listeria* infection drove NLRP6 activation. A hallmark of live cytosolic *L*isteria is the ability of the bacteria to induce the ActA-dependent polymerization of actin into characteristic actin tails that propel the bacteria forward and allow cell-to-cell spread ^4^. Since this causes a severe disturbance to the host cytoskeleton and endolysosomal system, we tested if cell-to-cell spread played a role in NLRP6 activation by infecting NLRP6-expressing GFP-ASC^tg^ cells with wild-type and ActA-deficient *Listeria.* Interestingly, we found that in comparison with cells infected with wild-type bacteria, cells infected with τι*actA* bacteria formed significantly less ASC specks (**Fig. 5A**), even though τι*actA*-infected cells contained on average higher numbers of intracellular bacteria (**Fig. S5A**). Of note, we specifically analyzed the infected cell population only, since the ActA-deficient mutant infects fewer cells, due to the loss of cell-to-cell spread. Since this result indicated an important role for actin-based motility in activating NLRP6, we treated cells with the actin polymerization inhibitor Latrunculin A after invasion and found that in addition to blocking bacterial actin tail formation, it also reduced *Listeria*-induced ASC speck formation, thus phenocopying *Listeria* ActA-deficiency (**Fig. 5A, S5B**). To test the importance of actin-based motility in a more physiological setting, we infected NLRP6 expressing HIEC-6 cells. ActA-deficient *Listeria* showed diminished inflammasome activation as evidenced by decreased GSDMD cleavage and cell death compared to cells infected with wild-type *Listeria*, confirming the results obtained in HEK293T cells (**Fig. 5B-C**). In addition to *Listeria*, the cytosolic bacterium *S. flexneri* is well-known to use actin-based motility to spread from cell to cell ^6^. Consistently, we found that *Shigella* infection also caused NLRP6-dependent cell death in infected HIEC-6 cells (**Fig. 5D**). However, the level of NLRP6-dependent death remained relatively low, potentially because we observed much lower levels of infected cells and actin-positive *Shigella* than *Listeria* at similar timepoints (**Fig. S4C)**.

**Figure 5.**
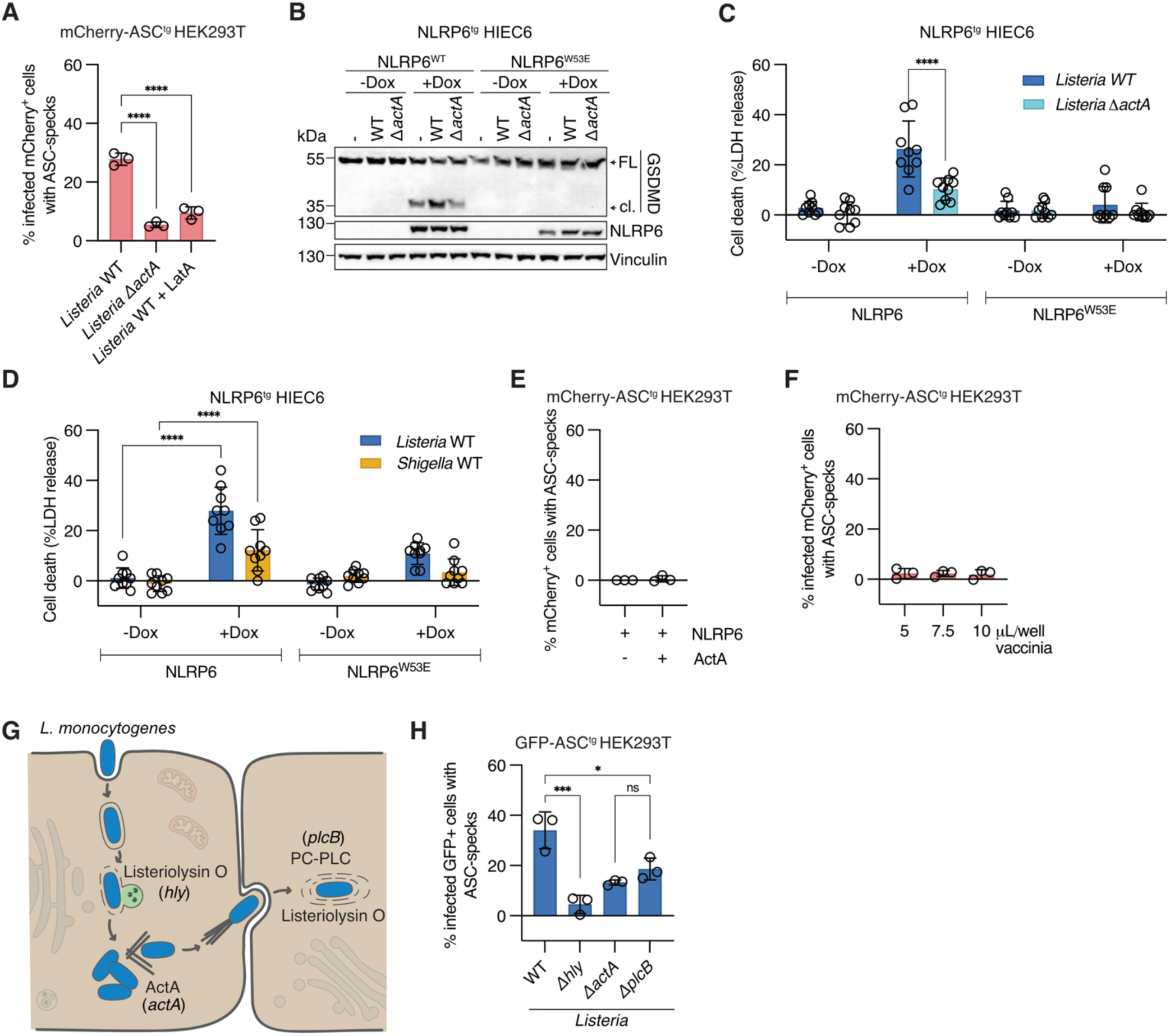
NLRP6 recognizes bacterial cell-to-cell spread. **A.** Flow cytometry-based quantification of ASC speck formation in hNLRP6-expressing mCherry-ASC^tg^ HEK293T cells, infected with GFP-expressing WT or τ<*actA L. monocytogenes* EGD for 6h, or infected with GFP-expressing WT *L. monocytogenes* EGD and treated with Latrunculin A (LatA) after 45min of infection. **B-C.** GSDMD processing and LDH release from NLRP6-WT^tg^ or NLRP6-W53E^tg^ HIEC-6 cells induced or not with doxycycline overnight and infected with WT or τ<*actA L. monocytogenes* EGD for 8h. **D.** LDH release from NLRP6-WT^tg^ or NLRP6-W53E^tg^ HIEC-6 cells induced or not with doxycycline overnight and infected with WT *L. monocytogenes* EGD or WT *Shigella flexneri* expressing AfaI for 8h. **E.** Flow cytometry based quantification of ASC speck formation in mCherry-ASC^tg^ HEK293T cells expressing hNLRP6 or hNLRP6 and ActA. **F.** Flow cytometry based quantification of ASC speck formation in hNLRP6-expressing mCherry-ASC^tg^ HEK293T cells, infected with indicated volume of GFP-expressing vaccinia virus at 4.7e10 pfu/mL for 8h. **G.** Schematic drawing showing *L. monocytogenes* virulence factors needed for primary vacuolar escape (Listeriolysin O, encoded by *hly* gene), actin-based motility (ActA, encoded by *actA* gene) and secondary vacuolar escape (PC-PLC, encoded by *plcB* gene and Listeriolysin O, encoded by *hly* gene). **H.** Flow cytometry-based quantification of ASC speck formation in hNLRP6-expressing mCherry-ASC^tg^ HEK293T cells, infected with WT, τ<*hly*, τ<*actA* and τ<*plcB L. monocytogenes* 10403S for 6h and stained for intracellular bacteria by immunofluorescence. Graphs A, E, F and H show mean ± SD from three pooled experiments. Graphs C and D show mean ± SD from three pooled experiments with three technical replicates each. Blots are representative of three independent experiments. FL (full length), cl. (cleaved). ns = non-significant, **P < 0.01; ****P < 0.0001 (one-way ANOVA with Dunnett’s multiple comparisons test (A) or Šídák’s multiple comparisons test (H), two-way ANOVA with Tukey’s multiple comparisons test (C) or Dunnett’s multiple comparisons test (D).

Finally, we tested if aberrant actin polymerization caused by the protein ActA itself triggered NLRP6 activation by ectopically expressing ActA in NLRP6-expressing GFP-ASC^tg^. Ectopically expressed ActA is reported to localize to mitochondria, where it nucleates actin ^43^. However, while ActA was expressed and did induce actin polymerization around mitochondria as expected, this did not induce NLRP6 activation (**Fig. 5E, S5D-E**). Furthermore, we infected NLRP6-expressing GFP-ASC^tg^ HEK293T cells with vaccinia virus that polymerizes host actin to promote spreading to new host cells ^6^. Vaccinia-infected cells did not show increased NLRP6 activation even though they were readily infected and featured actin-rich structures indicative of actin tails (**Fig. 5F, S5F-G**). Importantly, while these virus-induced actin tails appear similar to bacteria-induced actin tails, the actual spreading of vaccinia between cells is different, as the enveloped viral particle resides extracellularly at the tip of the actin tail, as opposed to bacterial spreading, where the bacteria remain intracellular and end up in a double membrane in the recipient cell ^44^. Altogether, these results suggest that it is not actin polymerization itself, but rather the bacterial cell-to-cell spread, which is dependent on actin polymerization, that is the driver of NLRP6 activation.

In contrast to the primary infection, which involves the escape from a single membrane endolysosome, actin-based invasion of neighboring cells (secondary infection) requires that the bacteria escape from a double membrane vacuole formed by plasma membranes of the cell of origin and the newly invaded cell. Since Listeriolysin O is insufficient to allow escape from such double membrane vacuoles, *Listeria* uses its phospholipase PC-PLC (encoded by the *plcB* gene) in combination with Listeriolysin O to destroy these membranes and enter the cytosol (**Fig. 5G**). Consequently, τι*plcB* bacteria are impaired in secondary vacuolar escape but not in primary invasion ^45^. To confirm that escape from double membrane vacuoles during secondary infection was necessary for NLRP6 activation, we compared NLRP6-expressing GFP-ASC^tg^ cells infected with wild-type, τι*hly*, τι*actA* and τι*plcB Listeria*, and found that τι*plcB Listeria* induced reduced levels of ASC-speck formation in the infected cell population, mimicking the effects of ActA deficiency (**Fig. 5H**). Consistent with published results, we found that PC-PLC deficiency (τι*plcB)* did not alter primary escape in the cells as seen by the number of infected cells and actin positive bacteria shortly after invasion but resulted in higher levels of LAMP1-positive (membrane bound) *Listeria* compared to the WT strain at later timepoints due to the failure of τι*plcB Listeria* to escape from double membrane vacuoles during secondary infection (**Fig. S4H-J**). These findings support the conclusion that escape from the double membrane vacuole during secondary infection of cells is necessary for complete NLRP6 activation by *Listeria*. It is however important to note that ActA and PC-PLC deficiency did not reduce NLRP6 activation to the level elicited by the Listeriolysin O-deficient mutant, which indicates that while escape from the secondary vacuole is the main driver of NLRP6 activation, another event, for example primary vacuolar escape, could still contribute to NLRP6 activation (**Fig. 5H**).

### Sterile endolysosomal damage is sufficient to activate NLRP6 in human intestinal cells

Since our findings had shown that secondary infections and the escape from double membrane vacuoles were crucial for NLRP6 activation, we wondered if NLRP6 specifically recognized the lysis of double membrane vacuoles over the lysis of single membrane vacuoles, or if the sheer number of escaping bacteria during late timepoints triggered NLRP6 activation. In support of the later hypothesis, NLRP6 activation was only detectable at 3-4 h post infection in both NLRP6^tg^ HIEC-6 and NLRP6 expressing GFP-ASC^tg^ HEK293T cells (**Fig. 1B, 2E**), at which time *Listeria* had already replicated around 10-fold (**Fig. S4E**). We thus asked if other triggers of lysosomal damage could activate NLRP6 if it occurred frequently enough. L-leucyl-L-leucine methyl ester (LLOMe) is a lysosomotropic agent that polymerizes inside lysosomes to induce rapid rupture of endolysosomal compartments and is thus often used to study the effects of vacuolar rupture, membrane repair and lysophagy ^46–48^. We first tested if LLOMe induces endolysosomal permeabilization in HIEC-6 cells by staining for Galectin-3, which binds β-galactosides on the luminal side of endolysosomes and thus can serve as a marker for permeabilization (**Fig. 6A**) ^49^. LLOMe caused endolysosomal permeabilization that was highest at 1 h post treatment and declined by 4 h, in line with published reports showing that damaged lysosomes are eventually either repaired or cleared (**Fig. 6A**) ^46,47,50–52^. Additionally, we also looked at endolysosomal membrane permeabilization during *Listeria* infection and found that while the wild-type bacteria induced Galectin-3 positive endolysosomal damage, the Listeriolysin O-deficient *τιhly* mutant did not (**Fig. 6A**). We next treated HIEC-6 cells expressing either wild-type or NLRP6^W53E^ with LLOMe and monitored inflammasome activation at 4 h post treatment. LLOMe treatment caused robust GSDMD cleavage in cells expressing WT NLRP6, in contrast to controls not treated with dox or expressing NLRP6^W53E^ (**Fig. 6B**). Furthermore, LLOMe induced lytic cell death in HIEC-6 cells expressing NLRP6 but not in the respective control cells (**Fig. 6C**). This indicates that while HIEC-6 cells are intrinsically resistant to LLOMe-induced lysosomal cell death, they could induce NLRP6-dependent pyroptosis in response to lysosomal disruption if they express NLRP6. Next, we asked if NLRP6-dependent pyroptosis induced upon LLOMe treatment required inflammatory Caspases. We thus treated NLRP6^tg^ HIEC-6 expressing sg*Luci*, sg*CASP1* or sg*CASP4* with LLOMe, and found that NLRP6-dependent pyroptosis triggered by LLOMe was exclusively dependent on Caspase-1 (**Fig 6D**). Since LLOMe caused the highest levels of endolysosomal damage at 1 h (**Fig. 6A**), we wondered if NLRP6 activation could be detected at these early timepoints. We thus measured PI uptake over time and found that loss of plasma membrane integrity could indeed be detected as early as 1-2 h post LLOMe treatment, thus coinciding with high levels of Galectin-3 staining (**Fig. 6E**). In addition to LLOMe, we also tested other triggers of endolysosomal damage, such as the dipeptide glycyl-L-phenylalanine 2-naphthylamide (GPN), which is a synthetic cathepsin C substrate, that upon cleavage induces endolysosomal disruption ^53^, as well as monosodium urate crystals, which once taken up into the cell will pierce and rupture endolysosomes ^54^. Treatment of HIEC-6 cells with GPN resulted in rapid NLRP6 dependent cell permeabilization and death (**Fig. 6F, S6A**), while treatment with MSU crystals induced only moderate levels of cell permeabilization, likely due to the non-phagocytic nature of HIEC-6 cells, but this permeabilization was also completely dependent on NLRP6 (**Fig. S6B**). Therefore, we could show that multiple triggers of sterile endolysosomal damage robustly and rapidly activate the NLRP6 inflammasome, thereby completely excluding a role for a bacterial PAMP in this activation.

**Figure 6.**
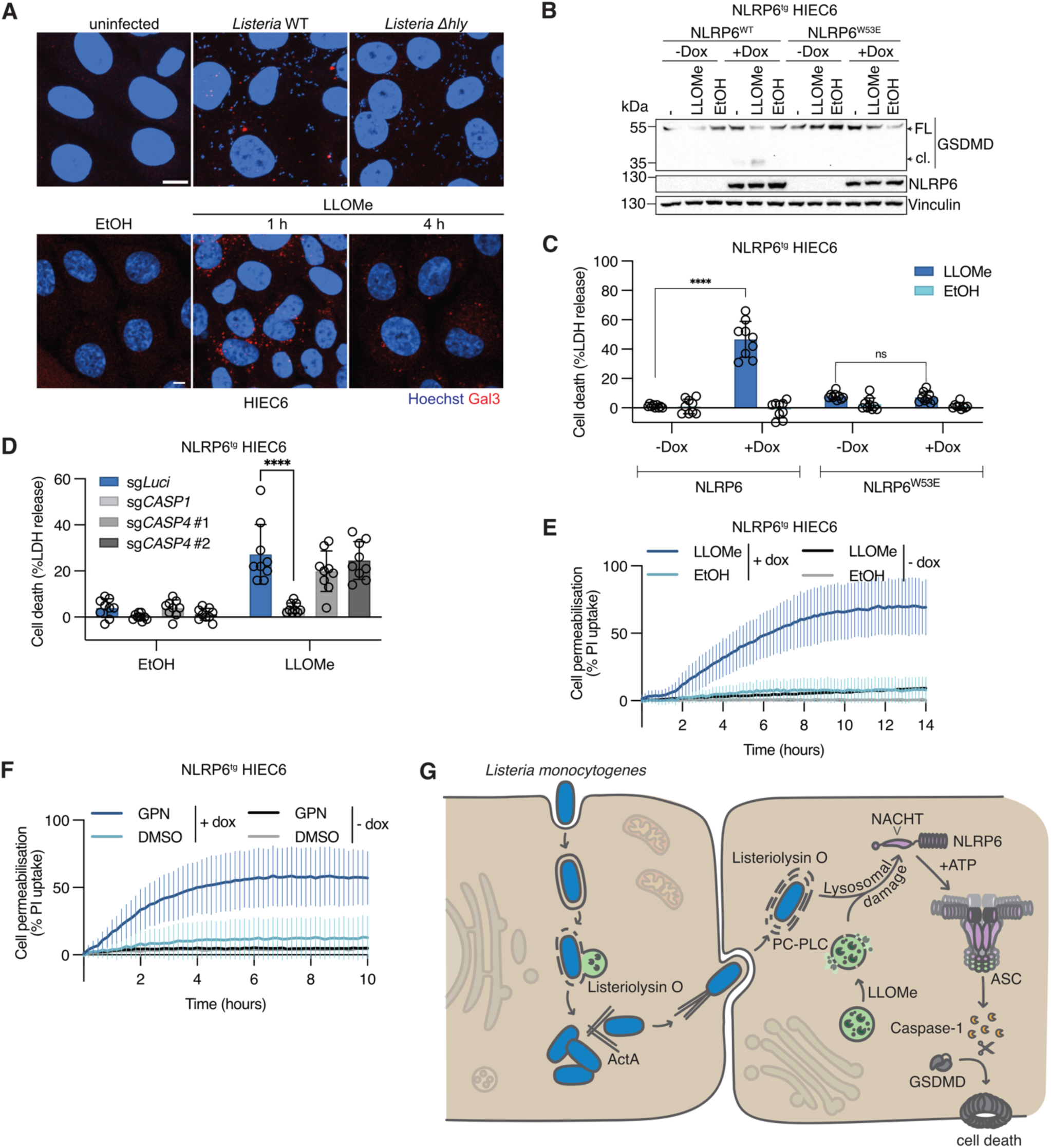
Sterile endolysosomal damage is sufficient to activate NLRP6 in human intestinal epithelial cells. **A.** Confocal maximum projection micrographs of HIEC-6 cells either infected with WT or τ<*hly L. monocytogenes* for 4h or treated with vehicle control (ethanol, EtOH) for 4h or 0.5mM LLOMe for 1h or 4h and stained for Galectin-3. **B-C.** GSDMD processing and LDH release from NLRP6-WT^tg^ or NLRP6-W53E^tg^ HIEC-6 cells induced or not with doxycycline overnight, left untreated or treated with 0.5mM LLOMe or an equivalent volume of ethanol (EtOH) as vehicle control. **D.** LDH release from NLRP6-WT^tg^ HIEC-6 controls cells (sg*Luci*) or polyclonal populations lacking Caspase-1 (sg*CASP1*) or Caspase-4 (sg*CASP4* #1 and #2) induced with doxycycline overnight and treated with 0.5mM LLOMe or equivalent volume of ethanol (EtOH) as vehicle control. **E-F.** Time course of propidium iodide (PI) uptake of NLRP6-WT^tg^ HIEC-6 cells induced or not with 1μg/ml doxycycline overnight and treated with 0.5mM LLOMe or equivalent volume of ethanol (EtOH) as vehicle control or treated with 200μM GPN or equivalent volume of DMSO as vehicle control. **G.** Summary scheme of proposed NLRP6 inflammasome activation by secondary spread of pathogenic bacteria or sterile lysosomal damage. Graphs C-F show mean ± SD from three pooled experiments with three technical replicates each. Images and blots are representative of three independent experiments. Scale bars represent 5μm. ns = non-significant, *P < 0.05, **P < 0.01, ***P < 0.001, ****P < 0.0001 (two-way ANOVA with Šídák’s multiple comparisons)

Endolysosomal damage results in ion (Ca^2+^) and proton fluxes, as well as the exposure of lipids and carbohydrates that can trigger the repair and regeneration of lysosomal membranes, or their removal by autophagy ^46,47^. Given the decline of Galectin-3 positive structures post LLOMe treatment we speculated that autophagy might contribute to NLRP6 activation. We thus first assessed the formation of autophagosomes in HIEC-6 cells treated with LLOMe and found increased size of LC3 puncta over time (**Fig. S5C-D**). We also found that LC3 puncta area increased over time in *Listeria* infected cells, consistent with reports that found an accumulation of pre-autophagosomal structures upon *L. monocytogenes* infection (**Fig. S5E-F**) ^55^. We next created NLRP6^tg^ HIEC-6 cells lacking ATG7, the E1 enzyme that is required for LC3 lipidation, a critical step in autophagosome biogenesis, and tested the functional impact of loss of autophagy on NLRP6 activation. As expected, when treated with LLOMe or infected with *Listeria*, ATG7-deficient cells did not show an increase of LC3B lipidation (LC3B-II), indicating a complete inhibition of autophagy (**Fig. S5G-H**). However, infecting these cells with *Listeria* did not cause a difference in NLRP6-dependent LDH release compared to the WT cells and NLRP6 activation by LLOMe treatment was only reduced in one of two guide RNA expressing populations, despite both gRNAs showing knockout and blockade of LC3 lipidation (**Fig. S5I-J**). Altogether, these results indicate that autophagy is not playing a role in NLRP6 activation.

In summary, we showed that NLRP6 is not exclusively detecting cell-to-cell spread of pathogenic bacteria and the resulting vacuolar damage but can also detect general endolysosomal damage caused by sterile triggers and thus acts as a general sensor for endosomal integrity independent of bacterial PAMPs.

## Discussion

Our study demonstrates that active NLRP6 nucleates an inflammasome and induces pyroptosis in response to both sterile and pathogen-induced lysosomal damage, thus establishing NLRP6 as a sensor of HAMPs (homeostasis-altering processes)^56^. Interestingly, we observe that during *Listeria* infection, NLRP6 activation is rarely triggered in primary infected cells as shown using mutant strains deficient for actin-based motility and the lysis of secondary vacuoles, but mainly activated in secondary infected cells once bacterial replication and the resulting endolysosomal damage has reached a threshold level. Such a threshold response could be necessary to avoid cell death caused by basal levels of endolysosomal damage, while maintaining the ability to respond to pathogens that replicate to high numbers within the epithelium. This might be especially important in barrier tissues like the intestinal epithelium, since excessive cell death responses could easily lead to a loss of barrier integrity. Consequently, the NLRP6 response appears to be specifically tailored towards pathogens that spread within the epithelium using actin-based motility as opposed to pathogens that replicate in an individually infected cell, such as *Salmonella*. Among the pathogens that use actin-based motility for cell-to cell spread are *Listeria* and *Shigella* that infect the intestinal epithelium, spotted fever group (SFG) *Rickettsia* that infect endothelial cells and *Burkholderia* spp. which infect, among others, the respiratory epithelium. The localized expression of NLRP6 in the intestinal epithelium thus correlates well with our observation that NLRP6 can detect both *Listeria* and *Shigella* in human IECs, and it will be interesting to investigate if other epithelia feature other immune sensors able to detect bacteria spreading between cells. Based on our data, we propose a model where the overwhelming spread of pathogens between cells, as well as pronounced lysosomal damage generates a yet-to-be-defined signal that results in a NACHT-domain and ATP-binding dependent activation and oligomerization of NLRP6. Active NLRP6 then recruits ASC to activate Caspase-1, which cleaves GSDMD, thereby permeabilizing the plasma membrane and inducing pyroptotic cell death (**Fig. 6G**). Therefore, NLRP6 acts as a guard of the integrity of the endolysosomal system in IECs.

An important question that remains to be answered is what ligand or signal is sensed by NLRP6 during endolysosomal damage. The finding that sterile endolysosomal damage by LLOMe, GPN and MSU is sufficient to activate NLRP6 in human IECs in the complete absence of bacteria and their associated PAMPs implies that NLRP6 is not activated by a specific bacterial effector, PAMP or activity, but by endolysosomal damage itself. Damage to endolysosomes has been shown to trigger several responses that depend on the severity of the damage and range from the engagement of repair mechanisms and regenerative pathways to the removal of damaged endolysosomes by autophagy. Recently it has been shown that imminent membrane damage results in the exposure of sphingomyelin on the cytosolic leaflet of stressed endolysosomal membranes and the recruitment of the sphingomyelin receptor TECPR1 that initiates non-canonical autophagy or conjugation of ATG8 to single membranes (CASM), i.e. the direct conjugation of LC3 to lipids in the lysosomal membrane ^57–59^. Initial lysosomal membrane instability caused by damage can be stabilized by plugging with stress granules ^60^ and small damage can be repaired by the engagement of ESCRT complexes recruited by calcium efflux from damaged endolysosomes ^47,50^ or a rapid ESCRT independent pathway involving phosphoinositide-initiated membrane tethering and lipid transport ^51,52^. Larger damage can lead to the recruitment of galectins (Gal-1, -3, -8 and -9), of which Gal8 coordinates the recruitment of the autophagy machinery for degradation ^61^. Lysophagy, the targeted degradation of terminally damaged lysosomes by autophagy, is triggered through Galectin-8 mediated recruitment of the adaptor protein NDP52 to initiate local autophagsosome biogenesis to sequester the damaged lysosome ^49,62,63^. New lysosomes can also be regenerated from damaged membranes in a process dependent on LIMP2 and ATG8-dependent recruitment of TBC1D15 and dynamin-2 ^64^. Finally, transcriptional reprogramming through the activation of the transcription factor TFEB upregulates the expression of lysosome biogenesis genes to initiate a program of lysosomal replacement ^65^. Many of these responses, for example galectin recruitment, autophagy and TFEB activation, have been shown to be linked and initiated in a coordinated manner ^65,66^. However, if any of these responses, the release of endolysosomal content or the presentation of a damage-associated ligand are linked to NLRP6 activation remains to be defined. The fact that ATG7 did not play a role in NLRP6 activation suggests that neither non-canonical autophagy nor lysophagy are important for NLRP6 inflammasome activation. However, it is possible that NLRP6 could bind another factor recruited to the damaged endolysosomes. Alternatively, NLRP6 could also sense processes that are initiated upon endolysosomal damage, such as the release of proteases and lipases from the damaged organelles, which has been shown to have diverse consequences, including the induction of cell death ^47,67^. Interestingly, it is well known that NLRP3 also reacts to endolysosomal damage, caused for example by LLOMe, pathogens or the uptake of particulate matter like uric acid crystals or alum. However, it is likely that the endolysosomal damage-derived signal that is recognized by NLRP6 and NLRP3 is different, since it has been shown that potassium efflux is a driver of NLRP3 activation under these conditions and we found no activation of NLRP6 by the potassium ionophore, nor did extracellular potassium supplementation inhibit NLRP6 activation. However, this does not exclude that inhibition of retrograde trafficking or Golgi dispersion, which activate NLRP3 ^68^, could serve as activators of NLRP6 as well.

The finding that NLRP6 functions as a sensor for disrupted homeostasis contradicts previous findings that suggest that NLRP6 detects specific PAMPs such as LTA, dsRNA or the bacterial metabolite taurine ^17,18,23^. Even though we could efficiently deliver dsRNA and LTA into cells, we were not able to detect any NLRP6 activation by these molecules, while *Listeria* triggered efficient NLRP6 activation in both HEK293T and HIEC-6 cells. A possible explanation could be the lack of a co-factor that is needed for dsRNA or LTA recognition by NLRP6, in analogy to dsRNA-triggered NLRP6-induced IFN signaling that relies on the co-factor DHX-15 ^14^. On the other hand, it is also possible that the transfection methods previously used to deliver dsRNA or LTA into cells caused endolysosomal damage and thus resulted in NLRP6 activation. Thus, future studies on NLRP6 activators will need to consider the disruption of cellular homeostasis as a possible driver of NLRP6 activation. A role for dsRNA and LTA in driving *Listeria*-induced NLRP6 activation can further be excluded as we have found that bacterial mutants that fail to spread between cells (e.g. τι*actA* or τι*plcB)* do not induce NLRP6 activation in infected cells, even though they are present at higher numbers within those cells than wild-type bacteria. Moreover, the fact that we were able to activate NLRP6 in the complete absence of any bacterial PAMP by the induction of sterile endolysosomal damage shows that PAMPs are not essential for the activation of NLRP6.

Our work also allowed us to perform mechanistic studies on NLRP6, which had previously been impossible due to the lack of a simple cell-based system for NLRP6 activation. These findings show that activation of NLRP6 was crucially dependent on the ability of the receptor to bind ATP, thereby suggesting that NLRP6 assembles into ATP hydrolysis-dependent wheel-like structures like NLRP3 or NLRC4 upon activation ^37–39,41^. While this appears to contradict the model that NLRP6 undergoes liquid-liquid-phase separation to activate ^23^, we cannot exclude that LLPS-dependent activation of NLRP6 also involves formation of wheel-shaped inflammasome complexes. Further support for the notion that NLRP6 forms complexes similarly to other NLRs comes from the fact that NLRP6 features a FISNA domain that displays sequence and structural similarity with the NLRP3-FISNA. In NLRP3 the FISNA domain was initially proposed to sense potassium efflux and recently Xiao et al. reported that the FISNA transitions from a partially disordered state to a folded state during receptor activation and that it is crucially required for receptor oligomerization ^32,33^. Our study shows that the FISNA domain is also important for NLRP6 activation, and most likely undergoes similar conformational changes during receptor activation. We were able to replace the entire PYD-linker-FISNA region of NLRP6 with the respective region of NLRP3 without changing ligand specificity. Moreover, replacing the NLRP3 NACHT with the NLRP6 NACHT conferred NLRP6-like signal specificity onto NLRP3, thus implying that for both NLRP6 and NLRP3 signal specificity lies within their respective NACHT domains. An alternative explanation for our findings would be that *Listeria* sensing lies within the PYD-linker-FISNA domains of both receptors and could therefore be interchanged non-specifically, while nigericin and imiquimod sensing could be uncoupled and sensed by the NACHT domain of NLRP3. Due to the lack of an NLRP6 only activator, we were unable to finitely determine whether sensing of imiquimod, nigericin and *Listeria* are all mediated by the same domain.

Finally, the finding that NLRP6 is not just a sensor for pathogen infection, but a general sensor for perturbed homeostasis of the endolysosomal system could potentially link NLRP6 to a plethora of different pathological processes. Increased lysosomal membrane permeabilization has been implicated in different diseases such as lysosomal storage diseases, neuro-degenerative diseases such as Parkinson’s and Alzheimer’s, as well as cancer ^69–72^. The link of NLRP6 with both neuroinflammation and inflammatory bowel diseases such as Crohn’s disease opens fascinating new research questions in the light of NLRP6 being a cellular homeostasis guard and its potential contribution to these malignancies ^20,21^. Our study is limited in the regard that we were only able to use NLRP6 reconstitution systems due to the lack of cell models with endogenous NLRP6 expression. However, our presentation of an activation mechanism and the domains being implicated in this process opens the field to explore more complex endogenous systems, such as *in vivo* and organoid models to explore the contribution of NLRP6 to infection and autoinflammation.

## Resource availability

Data and resources used in this study are available from the lead contact.

## Acknowledgements

We would like to thank Dr. Pascale Cossart, Dr. Marc Lecuit, Dr. Daniel Portnoy, Dr. Jan-Willem von Veening, Dr. Jost Enninga, Dr. Florian Schmidt and Dr. Fabio Martinon for providing bacterial or viral strains, as well as constructs. We would furthermore like to thank Dr. Ella Biewener Hartenian for her help with viral infections, Dr. Jakub Began for his help with LTA BODIPY labeling and Lukas Bissegger for support with cloning and cell culture. We would also like to acknowledge the support from the UNIL imaging, FACS and animal core facilities. Further thanks go to the rest of the current and former members of the Broz laboratory for constructive discussions and support. This work was supported by SNF Project funding (310030B_198005, 310030B_192523) to PB.

## Author contributions

A.B., E.M.B., L.L. and P.B. designed research. All authors performed research and/or analyzed data. A.B. and P.B. wrote the manuscript. All authors discussed results and commented on the manuscript.

## Declaration of interests

The authors declare no competing financial interests.

## Supplemental Figures

**Supplemental Figure 1.**
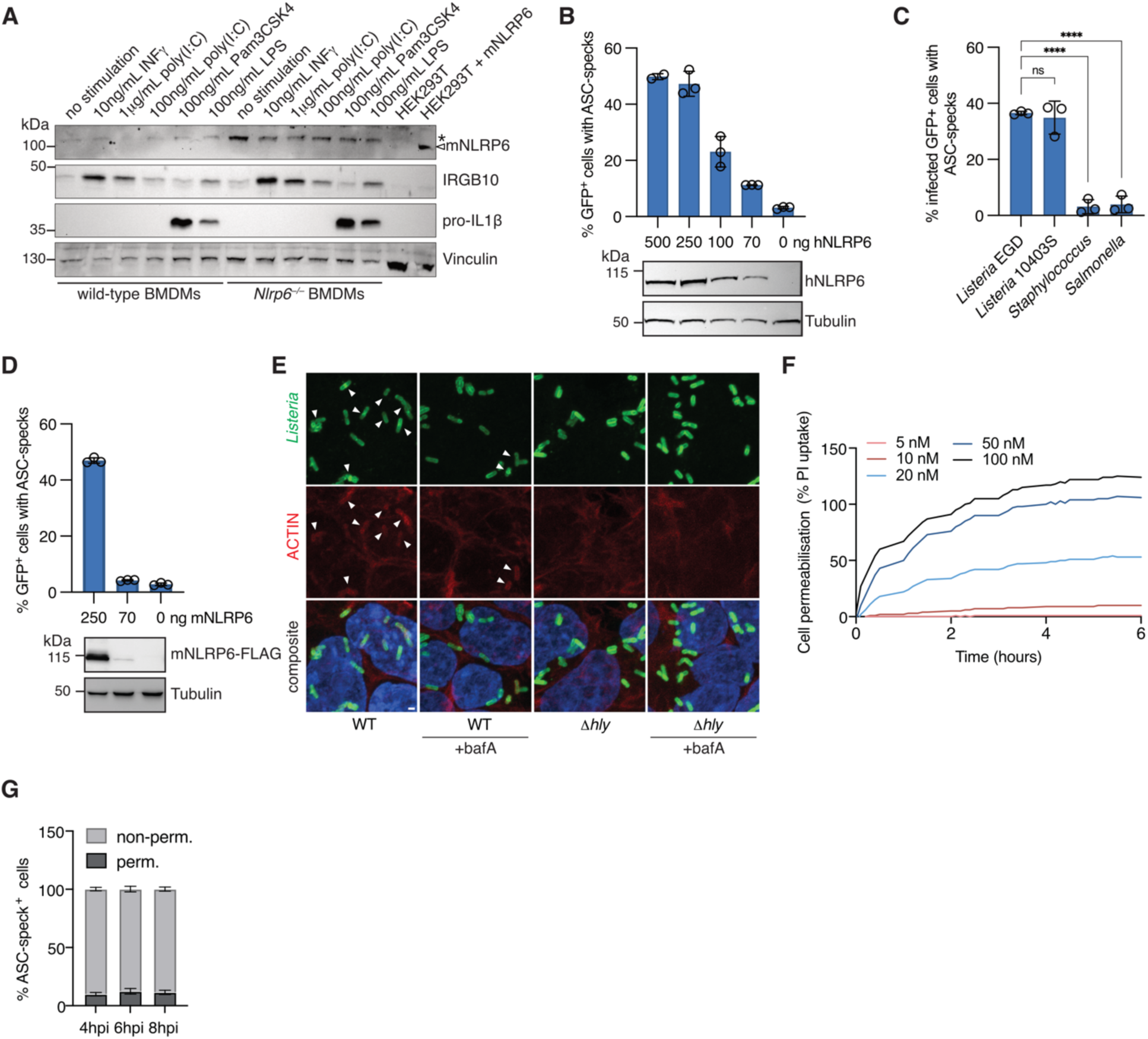
NLRP6 detects the entry of *L. monocytogenes* into the host cell cytosol. **A.** Immunoblotting for NLRP6 expression of murine wild-type or *Nlrp6^−/−^* BMDMs primed with 100ng/mL INFγ, 1μg/mL or 100ng/mL poly(I:C), 100ng/mL Pam3CSK4 or 100ng/mL LPS overnight. mNLRP6-FLAG expressing HEK293T cells are used as positive control. For successful priming, IGRB10 and pro-IL1β expression was used. Arrowhead indicates specific mNLRP6 band, while asterisk indicates non-specific band. Vinculin is used as a loading control. **B.** ASC speck formation quantified by flow cytometry and corresponding NLRP6 expression of GFP-ASC^tg^ HEK293T cells transfected with hNLRP6 DNA at indicated concentrations for 24h. **C.** ASC speck formation in hNLRP6-expressing GFP-ASC^tg^ HEK293T cells infected with *L. monocytogenes* strain EGD or 10403S*, S. aureus* or *S.* Typhimurium for 6h. **D.** ASC speck formation and corresponding NLRP6 expression of GFP-ASC^tg^ HEK293T cells transfected with mNLRP6 DNA at indicated concentrations for 24h **E.** Representative images of actin positive, intracellular *Listeria* after 1h of infection in HEK293T cells left untreated or treated with bafilomycin A (bafA) for 2h and infected with WT or τι*hly L. monocytogenes* EGD for 6h. *Listeria* were stained by immunofluorescene and Actin was stained with CellMask Actin Stain. Actin colocalization is indicated by arrowheads **F.** Propidium iodide (PI) uptake of HEK293T cells after treatment with purified Listeriolysin O at indicated concentrations over time **G.** Percentage of cells with permeabilized membranes among cells containing an ASC speck, hNLRP6-expressing GFP-ASC^tg^ HEK293T cells infected with *L. monocytogenes* EGD for 4, 6 or 8h as measured by flow cytometry LIVE/DEAD staining. Graphs show means ± SD from at three pooled independent experiments, except panel F, which is representative of two independent experiments. Images are representative of three independent experiments, immunoblots are representative of two independent experiments. Scale bar represents 1μm. ns = non-significant, ****P < 0.0001 (one-way ANOVA with Dunnett’s multiple comparisons).

**Supplemental Figure 2.**
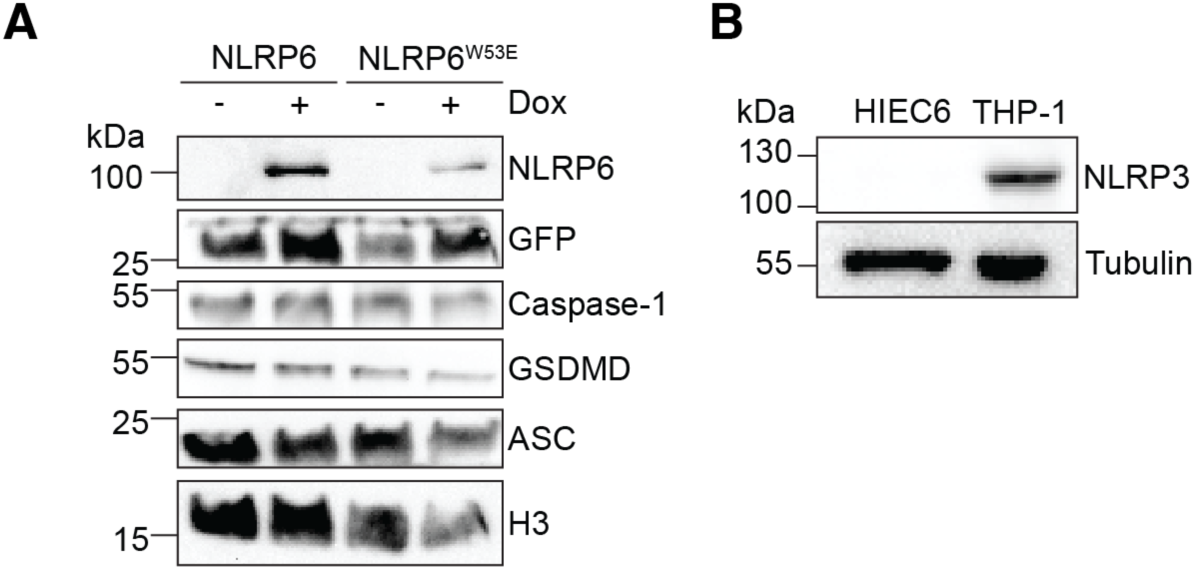
Expression of NLRP6 and other inflammasome components in HIEC-6 IECs. **A.** Immunoblots showing the expression NLRP6, GFP, Casp-1, GSDMD and ASC in NLRP6-WT^tg^ or NLRP6-W53E^tg^ HIEC-6 IECs left untreated or treated with 1μg/mL doxycycline overnight. Expression of GFP is shown as IRES-GFP was used as a marker to select the cells by flow cytometry. H3 serves as loading control. **B.** Immunoblots showing expression of NLRP3 in HIEC-6 IECs and THP-1 macrophages. Tubulin serves as loading control. Immunoblots are representative of three independent experiments.

**Supplemental Figure 3.**
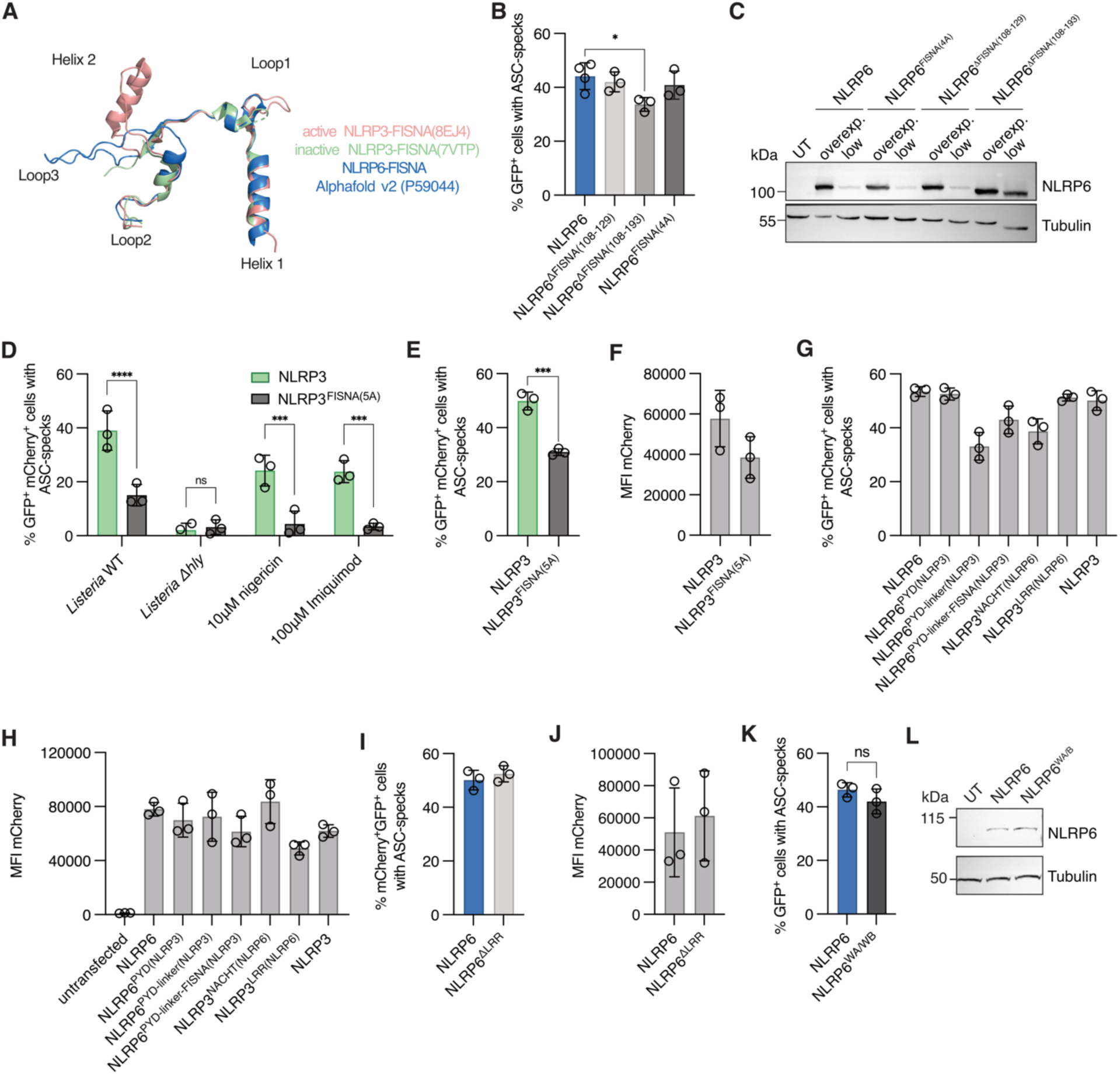
Molecular characterization of the NLRP6 receptor. **A.** Structural alignment of the FISNA domain of active and inactive NLRP3 (8EJ4 and 7VTP) with the NLRP6-FISNA predicted by Alphafold v2 (P59044). **B.** Flow cytometry-based quantification of ASC speck formation in GFP-ASC^tg^ HEK293T cells overexpressing the indicated NLRP6 proteins at 250ng DNA/well **C.** Immunoblot showing expression levels of constructs depicted in B expressed at either 250ng/well (overexp.) or 80ng/well (low). Tubulin serves as loading control. **D.** Flow cytometry based quantification of ASC speck formation in GFP-ASC^tg^ HEK293T cells, expressing the indicated NLRP3-mCherry proteins, infected with WT or τι*hly L. monocytogenes* for 6h or treated with 10μM nigericin or 100μM imiquimod for 1 or 6h respectively **E.** Flow cytometry based quantification of ASC speck formation in GFP-ASC^tg^ HEK293T cells overexpressing the indicated NLRP3-mCherry proteins at 250ng DNA/well **F.** Mean fluorescence intensity of NLRP3-mCherry in GFP-ASC^tg^ HEK293T cells expressing indicated NLRP3-mCherry proteins in D. **G.** Flow cytometry based quantification of ASC speck formation in GFP-ASC^tg^ HEK293T cells, overexpressing the indicated NLRP3-6-mCherry chimeric proteins at 250ng DNA/well. **H.** Mean fluorescence intensity of NLRP3-6-mCherry chimeric proteins in GFP-ASC^tg^ HEK293T cells transfected at 80ng DNA/well, except for chimeras NLRP3^NACHT(NLRP6)^ and NLRP3^LRR(NLRP6)^ which were transfected at 200 and 150ng/well each. **I.** Flow cytometry-based quantification of ASC speck formation in GFP-ASC^tg^ HEK293T cells, overexpressing the indicated NLRP6 proteins at 250ng DNA/well. **J.** Mean fluorescence intensity of indicated NLRP6-mCherry proteins in GFP-ASC^tg^ HEK293T cells transfected at 80ng DNA/well. **K.** Flow cytometry-based quantification of ASC speck formation in GFP-ASC^tg^ HEK293T cells overexpressing the indicated NLRP6 proteins at 250ng DNA/well **l.** Immunoblot showing expression levels of indicated NLRP6 proteins in GFP-ASC^tg^ HEK293T transfected at 80ng/well. Tubulin serves as loading control. Graphs show mean ± SD from three independent experiments. Immunoblots are representative of at least two independent experiments. ns = non-significant, *P < 0.05 ***P < 0.001, ****P < 0.0001 (one-way ANOVA with Dunett’s multiple comparisons test (B), two-way ANOVA with Šídák’s multiple comparisons test (D), unpaired t-test (E, K).

**Supplemental Figure 4.**
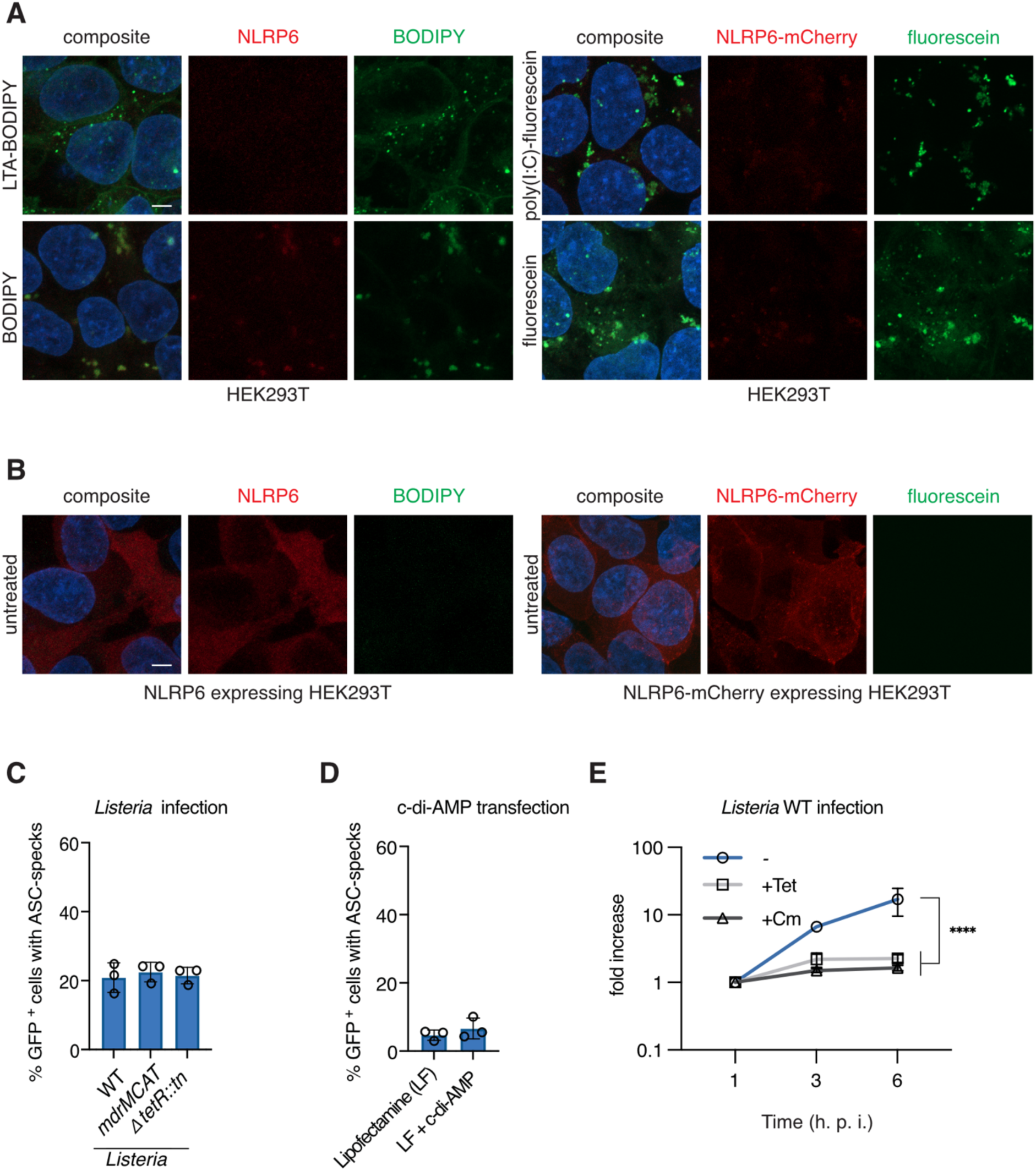
NLRP6 activation does not require bacterial PAMPs, but viable cytosolic bacteria. **A.** Representative maximum projection confocal micrographs after transfection of HEK293T cells with LTA-BODIPY (7.5μg/300,000), equivalent volume BODIPY only, poly(I:C)-fluorescein (250ng/300,000 cells) or equivalent volume fluorescein only for 6h. **B.** Representative maximum projection confocal micrographs hNLRP6- or hNLRP6-mCherry-expressing HEK293T cells left untreated. **C.** Flow cytometry based quantification of ASC speck formation in hNLRP6-expressing GFP-ASC^tg^ HEK293T cells infected with the indicated strains of *L. monocytogenes* 10403S for 8h. **D.** ASC speck formation in hNLRP6-expressing GFP-ASC^tg^ HEK293T cells mock treated or transfected with c-di-AMP for 24h. **E.** Replication of WT *L. monocytogenes* EGD over time in HEK293T cells in the presence of Tetracycline (Tet) or Chloramphenicol (Cm) added after 45min of infection. Graphs show means ± SD from at least three pooled experiments, except panel B, D and I, where two experiments were used. Images are representative of two independent experiments. Scale bars represent 5μm. ns = non-significant, ****P < 0.0001 (two-way ANOVA with Dunnett’s multiple comparisons test (E)).

**Supplemental Figure 5.**
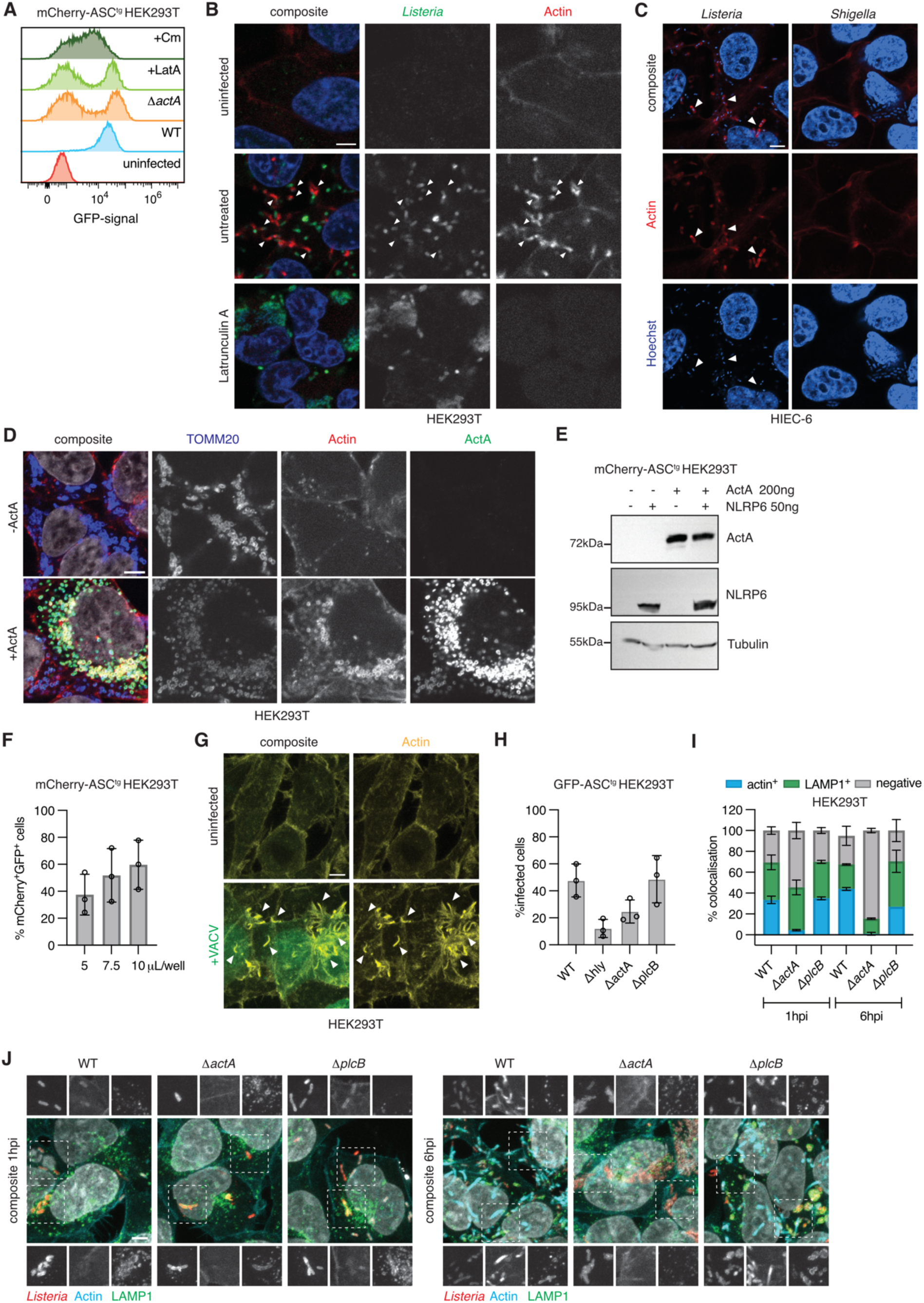
NLRP6 recognizes bacterial cell-to-cell spread. **A.** Distribution of GFP signal as analyzed by FACS in hNLRP6-expressing mCherry-ASC^tg^ HEK293T cells, infected with GFP-expressing WT or τι*act L. monocytogenes* EGD for 6h, or infected with GFP-expressing WT *L. monocytogenes* EGD and treated with Latrunculin A (LatA) or Chloramphenicol (Cm) after 45min of infection. **B.** Maximum projection confocal micrographs of actin tail formation by GFP-expressing WT *L. monocytogenes* EGD, treated with Latrunculin A (LatA) and imaged at 6hpi. Actin was stained with CellMask Actin Stain and actin tails are indicated by arrowheads. **C.** Maximum projection micrographs of actin tail formation by *L. monocytogenes* EGD or *S. flexneri* expressing AfaI after 6h of infection in HIEC-6 IECs, stained for Actin with CellMask Actin Stain. Arrowheads indicate actin-tails. **D.** Maximum projection confocal micrograph of HEK293T cells exogenously expressing ActA or not, stained for ActA and TOMM20 by immunofluorescence and Actin was stained with CellMask Actin Stain. **E.** Immunoblot for expression of hNLRP6 and ActA in mCherry-ASC^tg^ HEK293T cells expressing hNLRP6 or hNLRP6 and ActA. **F.** Percentage of vaccinia infected (GFP positive) hNLRP6-expressing mCherry-ASC^tg^ HEK293T cells, infected with indicated volumes of vaccinia virus at 4.7e10 pfu/mL for 8h. **G**. Actin tails formed in vaccinia virus infected (GFP positive) HEK293T cells. Actin was stained with CellMask Actin Stain and actin tails indicated by arrowheads. **H.** Percentage of infected cells in hNLRP6-expressing mCherry-ASC^tg^ HEK293T cells, infected with WT, τι*hly*, τι*actA* and τι*plcB L. monocytogenes* 10403S for 6h. Intracellular *Listeria* were stained by immunofluorescence and analysed by flow cytometry. **I-J.** Quantification and representative maximum projection confocal micrographs of intracellular *Listeria* colocalizing with Actin or the lysosomal marker LAMP1 in HEK293T cells infected with WT, τι*act* and τι*plcB L. monocytogenes* 10403S after 1 and 6hpi. Inserts are marked by dotted squares. *Listeria* and LAMP1 were stained by immunofluorescence and for Actin, CellMask Actin Stain was used. Graphs F and H show mean ± SD from three independent experiments. Graph I shows means ± SD from two independent experiments. Images and blots are representative of at least two independent experiments. Scale bars represent 5μm.

**Supplemental Figure 6.**
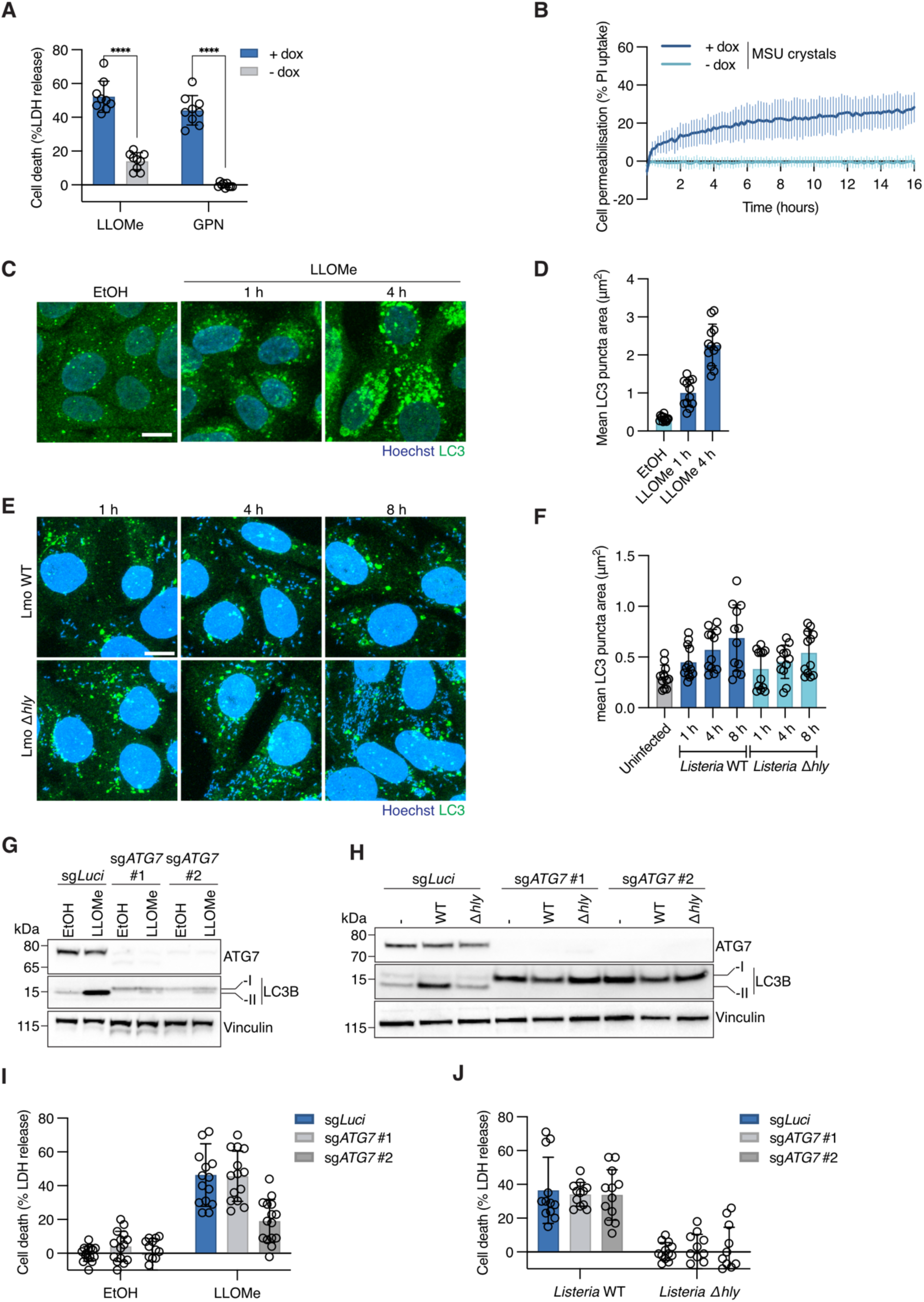
NLRP6 activation is independent of autophagy. **A.** Cell death measured by LDH release from NLRP6^tg^ HIEC-6 cells induced or not with 1μg/mL doxycycline overnight and treated with 0.5mM LLOMe or 200μM GPN for 8h. **B.** Propidium iodide (PI) uptake of NLRP6-WT^tg^ HIEC-6 cells induced or not with 1μg/ml doxycycline overnight and treated with 100μM MSU crystals for 15h **C-D.** Representative maximum projection confocal micrographs and quantification of LC3 puncta area of HIEC-6 cells treated with vehicle control (ethanol, EtOH) for 4h or 0.5mM LLOMe for 1h or 4h and stained for LC3 by immunofluorescence. **E-F.** Representative maximum projection confocal micrographs and quantification of LC3 puncta area of NLRP6^tg^ HIEC-6 cells induced with 1μg/ml doxycycline overnight and infected with *L. monocytogenes* EGD WT or τι*hly* mutant for 1, 4 or 8h and stained for LC3 by immunofluorescence. **G-H.** ATG7 expression and LC3 lipidation of NLRP6-WT^tg^ HIEC-6 controls cells (sg*Luci*) or polyclonal populations lacking ATG7 (sg*ATG7* #1 and #2) induced with 1μg/ml doxycycline overnight, either treated with 0.5mM LLOMe or an equivalent volume of ethanol (EtOH) for 4h or infected with *L. monocytogenes* EGD WT or τι*hly* mutant for 8h. Vinculin was used as a loading control. **I-J.** LDH release from NLRP6-WT^tg^ HIEC-6 controls cells (sg*Luci*) or polyclonal populations lacking ATG7 (sg*ATG7* #1 and #2) induced with 1μg/ml doxycycline overnight and treated with 0.5mM LLOMe or equivalent volume ethanol (EtOH) for 4h or infected with *L. monocytogenes* EGD WT or τι*hly* mutant for 8h. Graph A and B show mean ± SD of three or two independent experiments with three technical replicates each respectively. Graphs D and F show mean ± SD from two pooled experiments with 6 fields of view and at least 200 cells analyzed per condition and experiment. Each data point represents one field of view. Images and blots are representative of at least two independent experiments. Scale bars represent 5μm.

## Materials and Methods

### Bacterial strains and growth conditions

*Listeria monocytogenes* EGD wild-type and *τιhly* were a kind gift from Dr. Pascale Cossart (Institute Pasteur, France) and were grown in BHI medium (BD #237500) at 37°C ^73^*. L. monocytogenes* EGD τιactA mutant and corresponding wild-type, expressing either GFP or not, were a kind gift from Dr. Marc Lecuit (Institute Pasteur, France) ^74^. *L. monocytogenes* 10403S and all its mutants were grown in BHI medium supplemented with streptomycin (50µg/mL, Sigma-Aldrich #S9137) and were a kind gift from Dr. Daniel Portnoy (University of California, USA) ^42,45^. *Staphylococcus aureus* JE2 was grown in TSB (BD #211825) at 37°C and was a kind gift from Dr. Jan-Willem van Veening (University of Lausanne, Switzerland). *Salmonella enterica* serovar Typhimurium strain SL1344 was grown in LB supplemented with 10 g/L NaCl and streptomycin (50µg/mL) ^73^. *Shigella flexneri* M90T expressing the adhesin AfaI was a kind gift by Dr. Jost Enninga (Institut Pasteur, France) and was grown in TSB supplemented with ampicillin (50 μg/ml, Sigma-Aldrich #A9518) ^73^.

### Cell lines and Culture Conditions

HEK293T cells (ATCC CRL-3216) and BSC-40 (ATCC CRL-2761) were maintained in Dulbecco’s modified Eagle medium (DMEM) with GlutaMAX and pyruvate (Gibco #31966021), supplemented with 10% FCS at 37°C and 5% CO2. HIEC-6 cells (ATCC CRL-3266) and grown in Opti-MEM supplemented with 4% FCS, 10 mM Glutamine and 10 ng/mL of epidermal growth factor (EGF). WT and *Nlrp6^−/−^* mouse bone marrow-derived macrophages (BMDMs) from C57BL/6 male and female mice were harvested and differentiated in Dulbecco’s modified Eagle medium (DMEM) (Gibco) containing 20% L929 supernatant, as a source of macrophage colony-stimulating factor (M-CSF), 10% FCS (BioConcept), 10 mM HEPES (BioConcept), 1% penicillin/streptomycin and nonessential amino acids (NEAA, Gibco), and used for experiments on day 9 or 10 of differentiation.

### Vaccinia Virus Propagation

VACV Western Reserve EL EGFP was kindly provided by Dr. Florian Schmidt (University of Bonn, Germany). It was propagated on BSC-40 cells. Two 90% confluent T150 flasks (TPP #90151) were infected at MOI 0.01 and 2 days later scraped and subjected to two freeze thaw cycles. Cellular debris was centrifuged out at 300 x*g* for 10 minutes and the supernatant was centrifuged at 15 000 x*g* for 90 minutes. The viral pellet was resuspended in DMEM and titered on BSC-40 cells, after which single use aliquots of 4.7e10 pfu/mL were frozen at -80°C for infection ^75^.

### Animals

All experiments implicating animals (mice C57CL/6, male and female 8–12 weeks) were performed under the guidelines and approval from the Swiss animal protection law (licenses VD3257, Service des Affaires Vétérinaires, Direction Générale de l’Agriculture, de la Viticulture et des Affaires Vétérinaires, état de Vaud). All mice were bred and housed in a specific-pathogen-free facility at 22 ± 1 C° room temperature, 55 ± 10% humidity and a day/night cycle of 12 h/12 h at the University of Lausanne. *Nlrp6*-deficient mice have been described before ^22^.

### Construction of Plasmids and Lentiviral Transduction

All primers used in this study can be found in Table S1. ASC-mCherry was generated using an expression plasmid for ASC-GFP (Sino Biological #HG11175-ANG) as PCR template (PrimeSTAR MAX DNA Polymerase, Takara Bio #R045A) and was cloned into pLJM1_EGFP (addgene #19319) backbone, using AgeI-BstBI (New England Biolabs #R3552L and #R0519L) restriction digest and In-Fusion cloning (Takara Bio #6396650). In this pLJM1 backbone, both the CMV promoter was replaced by an EF1a promoter using SnaBI-NheI (New England Biolabs #R0130S and #R3131S) restriction digest and InFusion cloning, as well as the Puromycin resistance cassette was replaced by a Hygromycin cassette using BamHI-KpnI (New England Biolabs #R3136S and #R3142S) restriction digest and In-Fusion cloning. Human NLRP6 expression plasmid was obtained from Sino Biological (#HG21799-UT), murine NLRP6-FLAG expression plasmid was obtained from GenScript (#OMu17754). Human NLRP6 mutants for W53E, Walker A/B motives, FISNA deletion and FISNA(4A) mutant were generated by site-directed mutagenesis PCR amplification using In-Fusion cloning according to manufacturer’s instructions. The WalkerA/B mutant was cloned in two consecutive steps, first mutating motif A and then motif B. pINDUCER21 was a kind gift from Fabio Martinon (University of Lausanne, Switzerland) ^76^. The gene for human NLRP6 WT or W53E was amplified by PCR from vectors described above and cloned into pINDUCER21 using the BstBI-SpeI (New England Biolabs #R3133S) restriction sites and In-Fusion cloning. Human NLRP3-mCherry constructs are derived from a pEGFP-C2-NLRP3 plasmid (addgene #73955), while human NLRP6-mCherry constructs were derived from the plasmid mentioned above. The receptors and their chimeras were cloned into pLJM1_EGFP backbones using AgeI-BstBI restriction digest and In-Fusion cloning. NLRP6-mCherry under a EF1a promoter was cloned by using AgeI-BstBI restriction digest and In-Fusion cloning in the aforementioned pLJM1_EF1a_EGFP backbone with Puromycin resistance cassette. Successful cloning of all constructs was verified by Sanger sequencing.

To generate the stable ASC-GFP transgenic (tg) HEK293T (GFP-ASC^tg^ HEK293T) cell line, human ASC-GFP expression plasmid obtained from Sino Biological was linearized by XbaI restriction digest (New England Bioloabs #R0145S) and transfected into HEK293T cells using X-tremeGENE 9 DNA transfection reagent (Roche #XTG9-RO). After 24h, the cells were treated with 200ug/mL Hygromycin (Invivogen #ANT-HG-1) to select for genome insertion. The cells were sorted using flow cytometry into a low expressing population to avoid auto-aggregation.

For the generation of stable mCherry-ASC^tg^ HEK293T cell line, Lentiviral particles were produced from HEK293T cells. Briefly, 1×10^6^ cells were transfected with 1.25μg of plasmid, 1.25μg psPax2 and 0.25μg pVSV-G using TransIT LT1 transfection reagent (Mirus Bio #MIR2300). The following day media was exchanged for 4mL DMEM + 10% FCS + 1% BSA (Sigma-Aldrich #A9647). 72 h post transfection lentiviral particles were collected and filtered through a 0.45μm syringe filter (Sarstedt #83.1826). 1mL Virus-containing supernatant was supplemented with 4μg/ml Polybrene (Sigma-Aldrich #TR-1003-G) used to transduce HEK293T cells through spinfection at 500 x*g* for 60min. 24h after transduction, cells were treated with 200ug/mL Hygromycin to select for genome insertion. The cells were sorted using flow cytometry into a low expressing population to avoid auto-aggregation.

To generate HIEC-6 stable cell lines, lentiviral particles were produced from HEK293T cells. Briefly, 3×10^6^ cells were transfected with 1.9μg of pINDUCER21, 1.9μg psPax2 and 0.2μg pVSV-G using TransIT LT1 transfection reagent. The following day media was exchanged for 4mL DMEM + 10FBS + 1% BSA. 72h post transfection lentiviral particles were collected and filtered through a 0.45μm syringe filter. Virus-containing supernatant was supplemented with 4μg/mL Polybrene and all 4mL of supernatant were used to transduce HIEC-6 cells through spinfection at 500 x*g* for 90min. Cells were expanded and transduced cells were sorted by flow cytometry for GFP positive cells.

### Generation of CRISPR-Cas9 Knock-out Cell Lines

All single guide RNAs used in this study can be found in table S2. CRISPR-Cas9 Knock-out cell populations were generated using the LentiCRISPRv2 plasmid ^77^ expressing Cas9 from Streptococcus pyogenes and the tracrRNA previously published ^78^. The guide RNA sequences were designed using the CRISPick platform from the broad institute. Briefly, the vector was digested using BsmBI (New England Biolabs #R0739L) and ligated with the annealed oligonucleotides using T4 ligase system (New England Biolabs #M0202L). Plasmid sequence was verified using Sanger sequencing. Lentiviral particles were produced from HEK293T cells as described above. 1mL Virus-containing supernatant was supplemented with 4μg/mL polybrene and used to transduce HIEC-6 cells through spinfection at 500 x*g* for 90min. 24h after transduction, cells were treated with 7.5μg/mL Blasticidin (Invivogen #ant-bl-1) to select for genomic integration. Efficient knock-out was confirmed 7- and 10-days post transduction by immunoblot.

### Cell Transfection

HEK293T cells were either plated onto 24mm round glass coverslips (thickness 0.17+/-0.005mm, Biosystems Switzerland #0117640) coated first with poly-Lysine (0.01% poly-Lysine in water, Sigma-Aldrich #P8920), followed by collagen coating (5% collagen in PBS, Sigma-Aldrich #C4243), at a density of 7.5 × 10^4^ cells/well for microscopy or onto collagen-coated 24-well plates (TPP #92424) at a density of 7.5 × 10^4^ cells/well for flow cytometry or immunoblot samples 24 h before transfection. Cells were then transfected with one or two expression plasmids using XtremeGene 9 DNA transfection reagent for 24 h, according to the manufacturer’s instructions. Human and murine NLRP6, as well as human NLRP3 containing expression plasmids and all chimeric constructs, were transfected at 80ng/well in a 24-well plate, except for the NLRP3^NACHT(NLRP6)^ and the NLRP3^LRR(NLRP6)^ which were transfected at 200 and 150ng/well respectively.

### Cell Infection and Stimulation

*L. monocytogenes* grown overnight was diluted 1/15 in BHI and grown to mid-exponential phase (OD_600_ = 0.6-0.8). *S. aureus* grown overnight was diluted to OD_600_ = 0.4 in TSB and incubated for another hour to reach OD_600_ = 1. *S*. Typhimurium grown overnight was diluted 1/50 in LB and grown to late exponential phase (OD_600_ =1.3-1.5). *S. flexneri* grown overnight was diluted 1/100 in TSB supplemented with ampicillin (50 μg/ml) and grown to mid-exponential phase (OD_600_ = 0.4-0.6). Before infection, bacteria were washed and resuspended in either DMEM + 10% FCS or OptiMEM (Gibco #31985070), depending on the experimental setup*. L. monocytogenes, S. aureus*, *S.* Typhimurium and *S. flexneri* were added to confluent HEK293T cells at MOI 20, while *B. thailandensis* was used at MOI 100 if not indicated otherwise. Plates were then centrifuged for 5min at 300 x*g* and incubated for 45min (*L. monocytogenes, S. aureus, S*. Typhimurium and *S. flexneri*) at 37°C, 5% CO_2_. Noninternalized bacteria were eliminated using 20ug/mL Gentamicin (*L. monocytogenes*, Gibco #15710049), 100ug/mL Gentamicin (*S.* Typhimurium and *S. flexneri*) or 2ug/mL Lysostaphin (*S. aureus,* Sigma-Aldrich #L7386). For viral infection with vaccinia virus, cell medium was replaced for OptiMEM and the desired volume of viral suspension was added as indicated (titer 4.7e10 pfu/mL). The cells were incubated for one hour at 37°C, 5% CO_2_, after which the cell medium was replaced with DMEM + 10% FCS. At desired time points of infection, cells were processed for flow cytometry, Immunoblot, or microscopy. For CFU measurements, at desired time points, cells were washed with PBS, lysed with 0.2% Triton-X100 (AppliChem #A4975) in PBS (Gibco #14190-094), serially diluted and plated in duplicates on BHI agar. The agar plates were incubated for 24h at 37°C, after which colonies were counted.

For infection of HIEC-6 cells, 4×10^4^ cells/well were seeded in a 96-well plate (TPP #92696) and the following day, NLRP6 expression was induced overnight using 1μg/mL doxycycline (Sigma-Aldrich #D3447). Bacteria were cultured as stated above. Before infection, bacteria were washed and resuspended in OptiMEM. *L. monocytogenes* and *S. flexneri* were added at MOI 20. Plates were then centrifuged for 5min at 300 x*g* and incubated for one hour at 37°C, 5% CO_2_. Noninternalized bacteria were eliminated using 20μg/mL Gentamicin (*L. monocytogenes*) or 100μg/mL Gentamicin (*S. flexneri*). At desired time points of infection, cells and/or cell supernatants were processed for Immunoblot or cell permeabilization/death analysis. To induce endo-lysosomal membrane damage, NLRP6-expressing cells were treated with 0.5mM LLOMe (CHEM-IMPEX INT’L INC #04578), 200μM GPN (Abcam #ab145914), 100μM MSU (Invivogen #tlrl-msu) for indicated time points, after which cells and/or cell supernatants were processed for immunoblot or cell permeabilization/death analysis.

For ligand stimulation studies, unlabelled LTA (from *S. aureus*, Invivogen #tlrl-pslta) was transfected into cells at 10ug/300.000 cells, while BODIPY labelled LTA (labelling protocol below) was transfected at a theoretical concentration of 7.5ug/300.000 cells using DOTAP transfection reagent (Roche #11202375001). Glycine quenched BODIPY was transfected in equal volume. Briefly, the required volume LTA, LTA-BODIPY or BODIPY and 15uL DOTAP were separately diluted to 28uL in HBSS, mixed and incubated for five minutes, after which the LTA suspension was mixed with the diluted DOTAP and the mixture was brought to 200uL HBSS (Gibco #14025050). Of this mixture, 66.7uL/300.000 cells was added in each well on a 24-well or 96-well plate. For transfection with dsRNA, cells were transfected with 10ug/300.000 cells unlabelled LMW (Invivogen #tlrl-picw) or HMW poly(I:C) (Invivogen #tlrl-pic) or 0.25ug/300.000 cells for fluorescein-labelled HMW poly(I:C) (Invivogen #tlrl-picf) or glycine quenched fluorescein (Sigma Aldrich #46950) in equal volume using Lipofectamine 2000 (Thermo Fisher Scientific #11668019). Briefly, 1.5uL Lipofectamine and the required volume poly(I:C) were separately diluted to 25uL in OptiMEM and incubated for five minutes, after which the diluted poly(I:C) was added to the Lipofectamine suspension, mixed and incubated for 20min. Of this mixture, 25uL/300.000 cells was added for a 24- or 96-well plate. Transfection of c-di-AMP (Invivogen #tlrl-nacda) was performed at 10μg/300.000 cells using Lipofectamine 2000 as described above. *L. monocytogenes* lysate was prepared as previously published ^17^. Briefly, 10^9^ bacteria grown overnight in BHI broth were pelleted, washed in 1mL PBS, incubated with 100μg/mL lysozyme (Thermo Fisher Scientific #BP535-1), 5mM EDTA (Invitorgen #15575020) and protease inhibitor (Roche #11836170001) for 30 minutes at room temperature, after which the solution was sonicated and centrifuged for one minute at 21.000 x g. The supernatant was then used as bacterial lysate.

### Quantification of ASC-speck formation by flow cytometry

For flow cytometry analysis, supernatant and trypsinized cells were collected by centrifugation, washed in PBS and fixed in 4% PFA (Electron Microscopy Sciences #15710) for ten minutes. If indicated, cells were permeabilized using PBS + 0.1% BSA + 0.1% saponin (Sigma Aldrich #47036) and intracellularly stained for *L. monocytogenes* (abcam #ab35132), followed by labelling of secondary antibody coupled to fluorophore (either Alexa fluor 568, Invitrogen #A10042 or Alexa fluor 405 Invitrogen #A31556). When indicated, live/dead staining was performed using the LIVE/DEAD™ Fixable Violet Dead Cell Stain Kit for 405 nm excitation (Thermo Fisher Scientific #L34955) according to the manufacturers protocol.

For flow cytometry a CytoFLEX machine from Beckman Coulter was used. Cells were determined by forward and side scatter profile, 15,000 events were gated based on singlets (SSC-H vs. SSC-A). Cells were then gated for ASC-GFP or -mCherry expressors, and if applicable for NLR-receptor expression based on mCherry signal, live or dead cells based on the live/dead stain, or infected cells based on the intracellular staining (Alexa fluor 568 or 405) or GFP signal. ASC-speck formation was assessed in these gated cells by gating on GFP/mCherry-H vs. -A and selecting the shift towards higher signal according to previously published methods ^24^. At least 5,000 cells per condition and experiment were analyzed for ASC-speck formation. The raw ASC-speck formation percentages were normalized by subtracting the amount of auto-speck formation from the NLRP6/3 transfected cells that were untreated or uninfected.

### Microscopy, time-lapse imaging, and image analysis

For fluorescent microscopy of fixed samples, cells were fixed with 4% PFA for ten minutes. Fixed cells were permeabilized using PBS + 0.1% BSA + 0.1% saponin. Cells were then incubated 1:100 in permeabilization buffer with either anti-*L. monocytogenes*, anti-NLRP6 (AdipoGen #AG-20B-0046-C100), anti-ASC (Santa Cruz Biotechnology #SC-22514-R), anti-TOMM20 (abcam #AB56783), anti-ActA (Abnova #MAB8953), anti-LAMP1 (DSHB #H4A3-s), anti-LC3 (MBL International #PM036) or anti-Galectin-3 for one hour, washed three times with PBS and incubated with secondary antibody coupled to Alexa fluor-488, 568 or -647 at 2μg/mL (Invitrogen, 488: #A11008 and #A28175, 568: Invitrogen #A10042 and #A1103, 647: #A32795 and #A21235). DNA was stained using Hoechst (1:5000, Invitrogen #H3570), while actin was stained using CellMask^TM^ deep red actin tracking stain (Invitrogen #A57245) and samples coverslips were mounted in Vectashield mounting media (AdipoGen #VC-H-1900-L010). Samples were analysed with a LEICA DMI 6000b wide field microscope using a 40X/1.25-0.75 NA (oil) objective or a Zeiss LSM800 confocal laser scanning microscope using a 40X/1.3 NA (oil) or 63X/1.4 NA (oil) objective, if required acquiring Z-stacks with a 1-μm step size.

Images were analysed, quantified and processed using Fiji software. Below, procedure for quantification is described in more detail.

#### Actin-positive bacteria

For figure 1f total number of bacteria were enumerated from maximum projections of the *Listeria* image using background subtraction with a rolling ball of 10. A manual threshold was determined to select bacteria; the same threshold was used for all images in one experiment. Next, the Watershed function was used to separate touching objects. The Analyze Particles function was used to quantify the number of total bacteria using a size constraint of 80-Infinity pixels and circularity=0.30-1.00. Lastly, Actin positive bacteria were enumerated manually.

For figure S5j actin positive bacteria were quantified from a single middle plane of the Z-stack images. The total number of *Listeria* were enumerated with the same functions as described above. To identify actin positive bacteria, the mean pixel intensity of the actin stain was determined in the area defined by the bacterial mask. The mean pixel intensity to be considered actin positive was set using the highest values of the ActA-deficient *Listeria* at 6hpi.

#### ASC speck quantification

ASC specks were quantified from maximum projections of the Z-stack images. Total number of cells were enumerated by counting nuclei stained with Hoechst, using a Median Blur with a radius=7. A manual threshold was determined to select nuclei; the same threshold was used for all images in one experiment. The Fill Holes and Watershed functions were used to process a uniform mask and separate touching objects. The Analyze Particles function was used to quantify the number of nuclei using a size constraint of 30-Infinity. ASC specks were quantified by using a Median Blur with a radius=4 and setting a manual threshold; the same threshold was used for all images in one experiment. The Analyze Particles function was used to quantify the number of total ASC specks using a size constraint of 0.1-Infinity and circularity=0.70-1.00. ASC-specks per cell were enumerated by manually correcting for nuclei associated with multiple ASC specks.

#### LTA and poly(I:C) colocalization with NLRP6

For LTA-BODIPY, BODIPY, poly(I:C)-fluorescein and fluorescein colocalization with NLRP6(-mCherry), NLRP6 positive cells were segmented using a Gaussian Blur with sigma=7 and setting a manual threshold. Next, the mask was homogenized using the Fill Holes function. Within each region of interest (ROI) defined by NLRP6 positivity, particles of LTA or poly(I:C) were identified using a Gaussian Blur with sigma=2 and a manual threshold. To identify colocalization, the mean pixel intensity of the NLRP6 signal was determined in the area defined by the particle mask. The mean pixel intensity to be considered NLRP6 positive was set by manually checking the NLRP6 signal values for the particles. For each quantification, three independent experiments were used with each ten fields of view. At least 18 ROIs per condition and experiment were defined according to the manual NLRP6 positivity threshold, which could be comprised of multiple cells due to touching of the cells. At least 800 BODIPY or fluorescein positive speckles were counted and analyzed per condition and experiment.

#### LC3 Puncta Analysis

Maximum projections of LC3 image Z-stacks were processed using a background subtraction with a rolling ball of 50 and a Gaussian Blur with sigma=1. Afterwards, a manual threshold was determined to select LC3 puncta; the same threshold was used for all images in one experiment. Next, the Analyze Particles function was used to calculate the area of each punctum. The mean area of the puncta in each field of view was obtained from the ‘Summary’ of results.

#### LAMP1-positive bacteria

LAMP1 positive bacteria were quantified from maximum projections of the Z-stack images. The total number of *Listeria* were enumerated with the same functions as described above. To identify LAMP1-positive bacteria, the mean pixel intensity of the LAMP1 staining was determined in the area defined by the bacterial mask. The mean pixel intensity to be considered LAMP1-positive was set using the highest values of the ActA-deficient *Listeria* at 6hpi.

### Immunoblot Analysis

Cells were lysed in 24-well plates with 1X LDS sample buffer diluted in 1M Tris pH7.4 + 4% SDS. For GSDMD cleavage immunoblots, proteins were precipitated from cell supernatants using chloroform and methanol. 1 volume of MeOH and 0.3 volumes of chloroform were added to the supernatant, vortexed then centrifuged at 14000 x*g* at 4°C for 10 min. The upper, aqueous phase was discarded and 1.3 volumes of MeOH added before being vortexed and centrifuged at 14000 x *g* at 4°C for 10min. Supernatant was aspirated, and protein pellets allowed to airdry before being resuspended in the cell lysates from the corresponding wells. Samples were boiled at 95°C for 7 min then separated on 12% Tris-Gly SDS-PAGE gels and transferred to nitrocellulose membranes. Membranes were blocked in 5% milk in TBS-T then incubated with primary antibodies for human and murine NLRP6 (murine antibody purified from rabbit serum, inoculated with recombinant peptide, produced by YenZym), ASC (AdipoGen #AG-25B-0006), Caspase-1 (AdipoGen #AG202-0948-C103), Caspase-4 (Abcam #ab22687), GSDMD (Abcam full length: #ab210070, cleaved #ab215203), flag-tag (Sigma Aldrich #F3165), IRGB10 (kind gift from Dr. Jonathan Howard, Gulbenkian Institute of Science)^79^, pro-IL1β (R&D Systems #AF-401-NA), vinculin (Abcam #ab91459) and tubulin (HRP conjugated, Abcam #ab40742) overnight at 4°C (all antibodies at 1:1000, except murine NLRP6 at 1:2000 and tubulin-HRP at 1:5000). Membranes were washed in TBS-T before incubation with HRP conjugated secondary antibodies (SouthernBiotech #4030-05 and #1034-05) and finally developed using chemiluminescence (SuperSignal West Pico PLUS Chemiluminescent Substrate, Thermo Fisher Scientific #34580) on an iBright digital imaging system.

### Cell permeabilization and death Analysis

Cell membrane permeabilization was measured by PI uptake assay. After one hour of infection, cell medium was replaced by OptiMEM + 20ug/mL Gentamicin + 12.5ug/mL propidium iodide. The lysis control cells were lysed using 0.1% Triton-X100. 0.5mM LLOMe, equivalent volume ethanol, 200μM GPN or the equivalent volume DMSO or 100mM MSU was added in OptiMEM + 12.5ug/mL propidium iodide. Red fluorescence was measured using a Biotek 5 plate reader with Gen5 software at 37°C, 5% CO_2_, reading every ten minutes. Percentage of PI uptake was calculated using the formula:

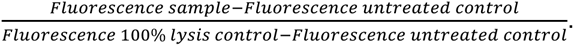

Lytic cell death was measured by LDH release assay (Sigma Aldrich #11644793001). 25μL of cell culture supernatant was incubated with 25μL of LDH assay reagent for 10-15 minutes before absorbance was measured at 490 nm on a Biotek plate reader. 100% lysis control was prepared by adding a final concentration of 0.1% Triton-X to the corresponding wells. All conditions were performed in technical triplicate and the percentage of LDH

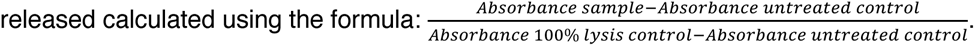

### Protein Sequence alignments

Protein sequences for NLRP6 and NLRP3 from different species were obtained from UniProt (UniProtKB accession numbers Q8R4B8, Q96P20, P59044 and Q91WS2) and a multiple protein sequence alignment was generated using Clustal Omega (www.ebi.ac.uk). Domains were identified and delineated based on previous publications ^23,25,32^ or homology to NLRP3.

### LTA labelling with BODIPY

Labelling protocol for LTA was adapted from previous publication ^17^. LTA was prepared as 5mg/mL stock in endotoxin free water. 500uL of this stock was diluted to 1.25mL and sonicated for ten minutes at 70% power on ice. After, 1.25mL 200mM Sodium Bicarbonate was added to a final LTA concentration of 1mg/mL (125uM). BDP FL NHS Ester (TCI, D5555) was added gradually to reach molar ratio of LTA to dye of 1:4. The solution was allowed to react for 1.5 hours at room temperature in the dark, after which the mixture was centrifuged at 15.000 rpm for 10 minutes to remove precipitants. The staining reaction was quenched with 2.5mM Glycine (Sigma Aldrich #G7126) at neutral pH for 30 minutes. Gel filtration of the quenched solution was performed on a Superdex 200 column and collected in the void fractions. From the final volume collected and pooled after gel filtration, concentration of labelled LTA was calculated to be 100μM (0.8mg/mL). The fractions containing the quenched BODIPY was used as a negative control for transfection of cells with fluorophore only.

#### Quantification and Statistical Analysis

Data analysis was performed using Flow Jo 10, GraphPad Prism v10, Microsoft Excel, and Fiji software. Statistical significance is indicated as non-significant (ns), *, **, *** or **** for a P value of >0.05, <0.05, <0.01, <0.001 or <0.0001, respectively. For comparison of two individual groups, a two-tailed, unpaired t-test was used, while for comparison of three or more groups, one-way ANOVA or two-way ANOVA was used with Dunnett’s/Tukey’s/ Šídák’s/Uncorrected Fisher’s LSD multiple comparisons test as indicated. Number of independent biological replicates is indicated in given figure legend, as well as number of events quantified for microscopy analysis.

**Table S1.**
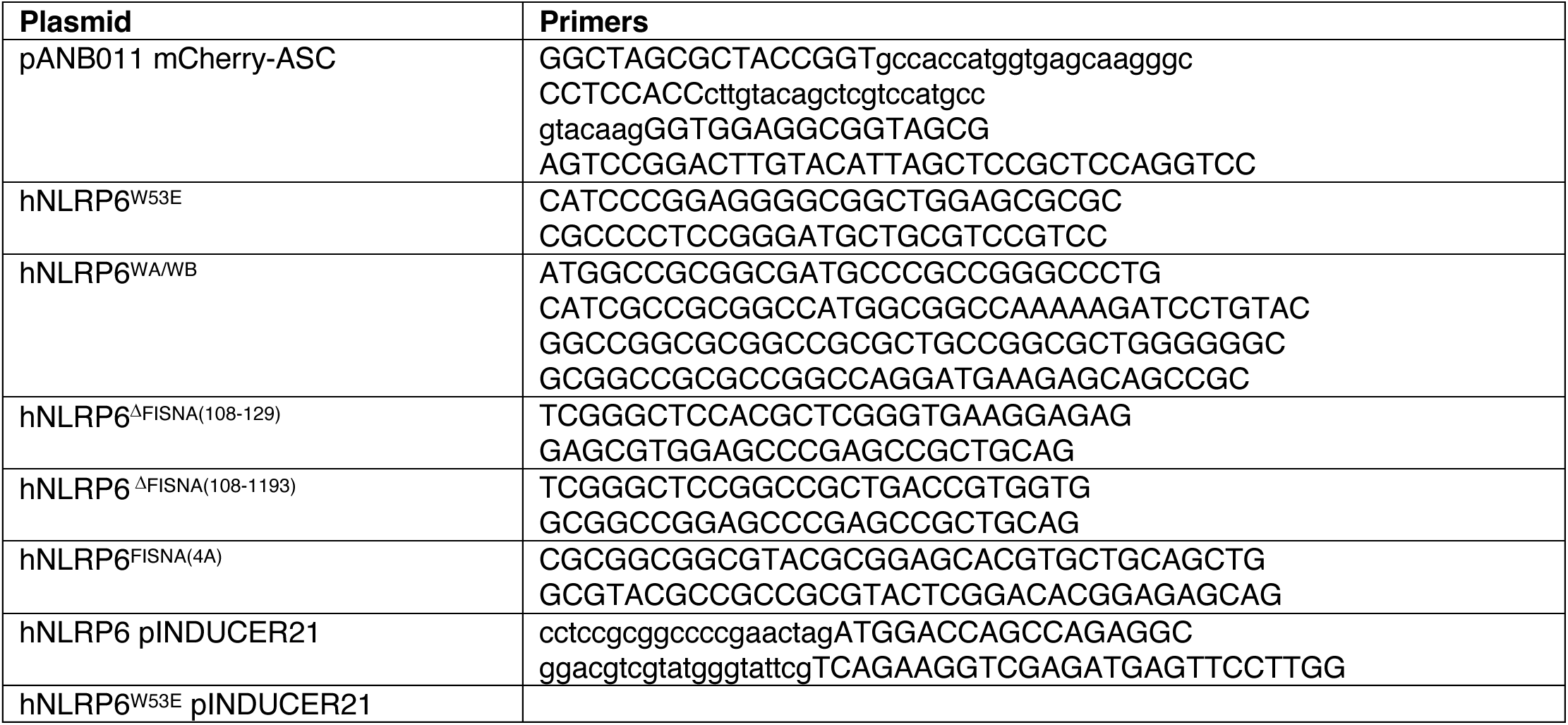
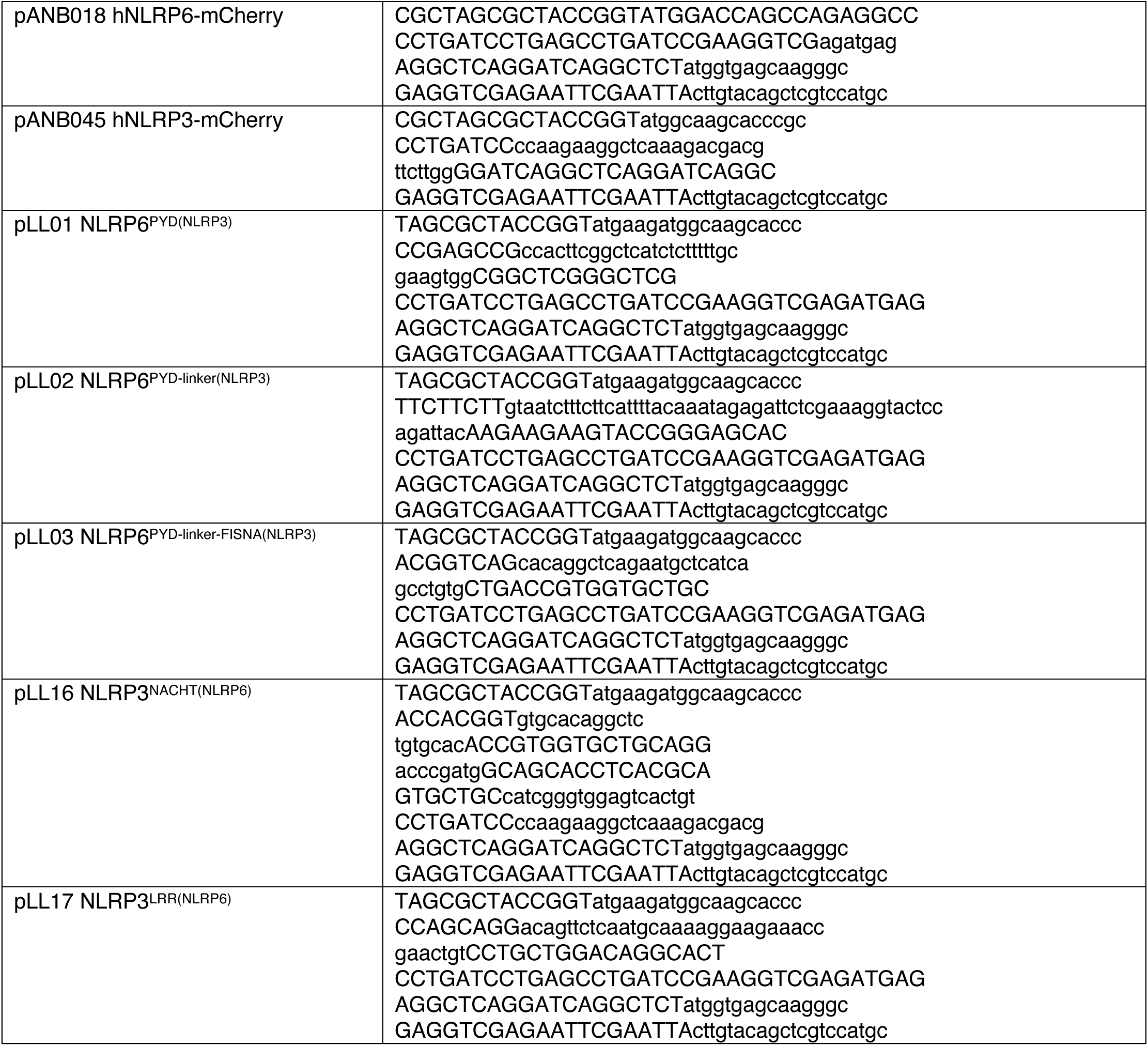
Primers used in this study to clone indicated plasmids.

**Table S2.**
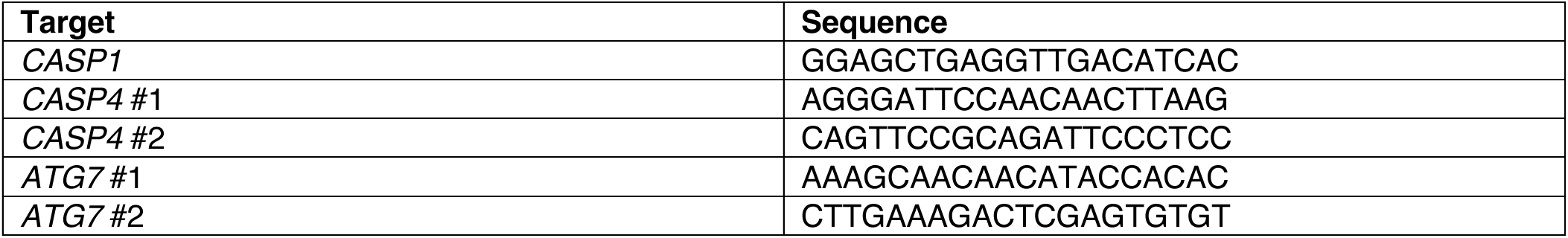
Single guide RNAs used in this study to create population wide knock-out using CRISPR-Cas9.

## Notes

### Competing Interest Statement

The authors have declared no competing interest.

